# Apomixis frequency in polyploid wild populations of *Psidium cattleyanum* f. *lucidum* (Myrteae, Myrtaceae)

**DOI:** 10.1101/2025.08.18.670884

**Authors:** M. Souza-Pérez, R Machado, M. Vaio, A. Borges, J. I. Hormaza, G. Speroni

**Affiliations:** Departamento de Biología Vegetal, Facultad de Agronomía, Universidad de la República Montevideo, Uruguay; Universidade Federal do Maranhão - Centro de Ciências de Grajaú, Av. Aurila Maria dos Santos Barros Sousa, 2010, 65940000, Grajaú, Maranhão, Brazil; Departamento de Biometría, estadística y computación, Facultad de Agronomía, Universidad de la República Montevideo, Uruguay; Instituto de Hortofruticultura Subtropical y Mediterránea la Mayora (IHSM La Mayora – CSIC – UMA), 29750 Algarrobo_Costa Málaga, Spain

**Keywords:** Ploidy levels, gametophytic apomixis, seed progeny, seeds-flow cytometry, reproductive mode, Embryo/Endosperm DNA content balance, unreduced gametes, *Psidium cattleyanum*, SSR genotyping

## Abstract

**Background and Aims:** Apomictic reproduction and polyploidy are closely related in plants, and their causes and consequences have given rise to numerous hypotheses that continue to evolve as more representative taxa are studied. *Psidium cattleyanum* is a valuable model for investigating this relationship, as it is a polyploid species with multiple ploidy levels and exhibits pseudogamous gametophytic apomixis. In addition, it is a fruit tree of commercial, agricultural, and medicinal interest in its native range, but it is also an aggressive invasive species in regions where it has been introduced. In this study, we aim to determine the predominant reproductive mode in natural populations of the polyploid *Psidium cattleyanum* f. *lucidum* and to investigate whether this mode is associated with specific ploidy levels. Furthermore, we investigate whether these reproductive traits are linked to patterns of genetic diversity and population genetic structure.

**Methods:** We assessed the reproductive mode in wild populations representing the four ploidy levels (2C = 5x, 6x, 7x, 8x) present in Uruguay. Reproductive pathways were inferred by combining flow cytometric seed screen (FCSS) analysis with microsatellite (SSR) genotyping of progeny arrays. Population-level differences were evaluated across both data sets.

**Key Results:** Pseudogamous apomixis is the predominant reproductive mode in Uruguayan *P. cattleyanum* f. *lucidum* populations. The proportions of apomictic versus sexual reproduction, and the types of progenies produced, differed significantly among populations and ploidy levels. Genotyping revealed predominantly clonal progeny, yet with high genetic variability, and populations exhibit strong genetic structure. The ploidy level of the gametes and their combinations were characteristic of each population, leading to different reproductive strategies.

**Conclusions:** Facultative pseudogamous apomixis is the primary reproductive mode in natural populations of *Psidium cattleyanum* f. *lucidum*. Ploidy level influences reproductive strategy, as reflected in the apomixis/sexuality ratios and the type of progeny produced. Population genetic structure is shaped by the reproductive strategy adopted by each population.

## Introduction

Apomixis, asexual reproduction via seeds, is widely distributed across the phylogeny of the angiosperms (Hörandl *et al*., 2024). However, approximately 75% of the well-documented cases are concentrated in just three of 74 families: Asteraceae, Rosaceae, and Poaceae (Hojsgaard and Pullaiah, 2022). The most common form among angiosperms is gametophytic apomixis (Hojsgaard and Pullaiah, 2022), which involves the formation of unreduced embryo sacs (ES) and embryo development through parthenogenesis of the egg cell. This process is termed diplospory if the embryo sac originates from the megaspore mother cell, and apospory when it originates from somatic cells in the ovule tissues (Nogler, 1984). Most gametophytic apomictic plants are pseudogamous, which means that the fertilization of the central cell of the embryo sac is required to form the endosperm (Mogie, 1992; van Dijk and van Damme, 2000), whereas a minority are autonomous apomictics, with the endosperm originating directly from the central cell without fertilization (Nogler, 1984).

Plant taxa that reproduce exclusively through apomixis are referred to as obligate apomictic. This reproductive strategy has been reported in several genera of Orchidaceae (Sorensen *et al*., 2009; Kant and Verma, 2012; Zhang and Gao, 2018; Xiao *et al*., 2021) and in species of other families, such as *Paspalum malacophyllum* (Poaceae) (Hojsgaard *et al*., 2013), *Arnica cordifolia* (Asteraceae) (Kao, 2007), and *Miconia albicans* (Melastomataceae) (Caetano *et al*., 2013). When apomixis coexists with sexual reproduction within the same taxon, it is described as facultative apomixis (Maheshwari, 1950), a condition thought to predominate among species exhibiting gametophytic apomixis (Hojsgaard and Pullaiah, 2022). Nevertheless, it is assumed that most apomictic species retain some capacity to reproduce sexually (Hojsgaard and Hörandl, 2015; Hodač *et al*., 2019; Hofstatter and Lahr, 2019). Moreover, the reproductive mode adopted by a plant may shift in response to environmental or developmental conditions, a process governed by genetic or epigenetic regulation (reviewed by Hörandl *et al*., 2024).

The majority of known apomictic angiosperms are polyploid (Asker and Jerling, 1992; León-Martínez and Vielle-Calzada, 2019; Hörandl *et al*., 2024). Particularly, a strong association exists between gametophytic apomixis and polyploidy. In several polyploid species and species complexes, diploid cytotypes typically reproduce sexually, while apomixis is restricted to polyploids (e.g*. Crataegus*, Lo *et al*., 2009; *Rubus*, Šarhanová *et al*., 2012; *Amelanchier*, Burgess *et al*., 2014; *Taraxacum*, Mártonfiová, 2015; *Pilosella*, Rotreklová and Krahulcová, 2016; *Handroanthus ochraceus*, Mendes *et al*., 2018; *Sorbus*, Lepší *et al*., 2019; *Hieracium*, Mráz and Zdvořák, 2019; *Cotoneaster integerrimus*, Bogunić *et al*., 2021; *Ranunculus kuepferi*, Ladinig *et al*., 2024). One of the primary mechanisms proposed to explain the apomixis-poliploidy correspondence is the formation of unreduced gametes, which are capable of either fertilizing or being fertilized, thereby contributing to polyploidization (Whitton *et al*., 2008; Hörandl *et al*., 2024). However, as this is not the sole driving force, for each plant group it is crucial to examine their preceding evolutionary and reproductive mechanisms to elucidate the underlying basis of this phenomenon.

Advances in techniques such as flow cytometric seed screening (FCSS) have significantly enhanced the ability to determine reproductive modes in plant species (Matzk *et al*., 2000). FCSS enables rapid and efficient analysis of large numbers of mature seeds by quantifying the nuclear DNA content, allowing the determination of the ploidy levels of both the embryo and the endosperm cells. From the resulting embryo:endosperm DNA content ratios, it is possible to infer the mode of reproduction, by assessing deviations from the double fertilization process typical of angiosperms (Krahulcová and Rotreklová, 2010) and to interpret the diversity of possible combinations taking into account the ploidy level of the gametophytes involved (Krahulcová *et al*., 2004; Talent and Dickinson, 2007; Qu *et al*., 2010; Šarhanová *et al*., 2012; Dobeš *et al*., 2013b; Lepší *et al*., 2019; Da Luz-Graña *et al*., 2025). In a *Polygonum*-type embryo sac (ES), the expected ploidy ratio under sexual reproduction is 2:3. In contrast, gametophytic apomixis typically yields a ratio of 2:4 under autonomous endosperm development and 2:5 under pseudogamous condition (Fig. 1). Other ratios may arise depending on whether the contributing sperm nuclei and the embryo sac are reduced or unreduced (Fig. 1). Accurate interpretation of FCSS data requires a detailed understanding of the embryological features of the species being investigated (Dobeš *et al*., 2013a; Hörandl *et al*., 2024).

**Figure 1:**
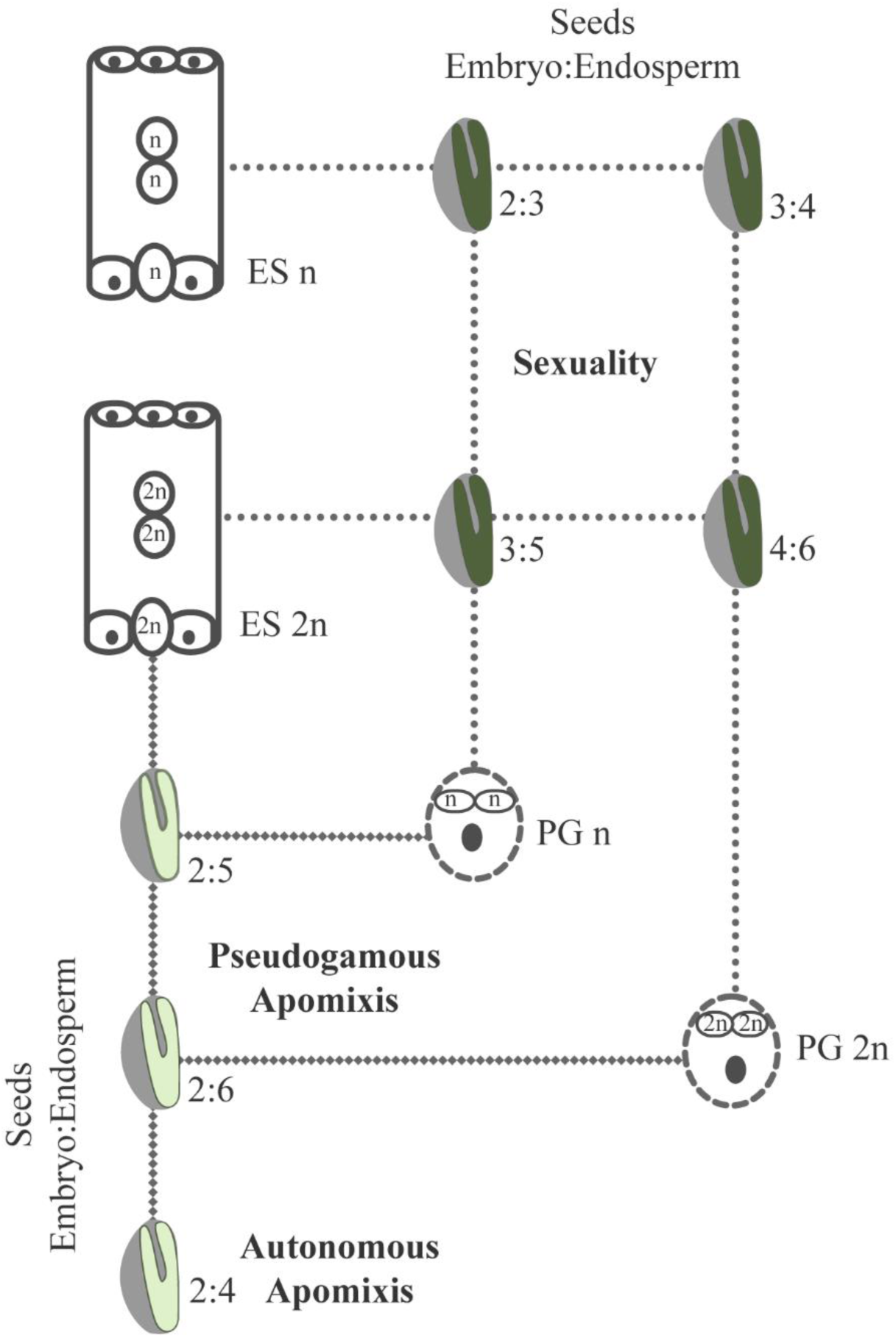
Reproductive modes inferred from seed scanning via flow cytometric seed screening (FCSS). Diagram based on the most common reproductive modes in angiosperms, derived from a Polygonum-type embryo sac and bicellular pollen grains. In sexual reproduction, one sperm nucleus fertilizes the egg cell and the other fertilizes the central cell (2:3, 3:5, 3:4, 4:6). In apomictic cases, the egg cell forms an embryo through parthenogenesis, and the endosperm is either autonomous, generated from the central cell nuclei (2:4) or pseudogamous resulting from fertilization of the central cell by a sperm nucleus (2:5). ES = embryo sac, PG = pollen grain.

Another widely used approach for determining the mode of reproduction involves genotyping of parents and offspring using molecular markers, such as microsatellites (SSRs) (Šarhanová *et al*., 2017, 2024; Hojsgaard and Pullaiah, 2022). SSRs are especially valuable for studying polyploids, as they are codominant markers and allow the detection of a high number of alleles (Vieira *et al*., 2016; Hodač *et al*., 2019). This analysis enables the assessment of genetic variability within facultatively apomictic populations and their progeny and complements FCSS, especially in cases where the method lacks resolution— for example, in autonomous endosperm formation, absence of central cell nuclei fusion before fertilization, instances of gametic aneuploidy or reduced/absent endosperm, and secondary chemical compounds interfering with DNA staining (Matzk *et al*., 2000; Talent and Dickinson, 2007; Krahulcová and Rotreklová, 2010; Jedrzejczyk and Sliwinska, 2010; Dobeš *et al*., 2013a).

*Psidium cattleyanum* is naturally distributed in South America along the Atlantic coast, ranging from Espírito Santo in Brazil to northeastern Uruguay (Sobral *et al*., 2006; Brussa and Grela, 2007). It is cultivated in several other countries and has become an aggressive invasive species in several tropical regions (Global Invasive Species Database, 2025). The species has significant agronomic potential due to high fruit yield and the presence of bioactive compounds in leaf and fruit extracts (Chaves *et al*., 2018; dos Santos Pereira *et al*., 2018; Zandoná *et al*., 2020). Furthermore, it is considered a valuable source of germplasm for the genetic improvement of the widely cultivated guava, *P. guajava* (De Almeida *et al*., 2012; Marques *et al*., 2012; Cardoso *et al*., 2017; Fortuna Macan and Cardoso, 2020).

*Psidium cattleyanum* is considered an obligate polyploid species, as no diploid cytotypes (2n = 22) have been reported to date (Proença *et al*., 2022). Multiple ploidy levels have been documented, including 2n = 44, 55, 66, 77, 82, 88, 99, 110, and 132 (Atchison, 1947; Costa and Forni-Martins, 2007; De Souza *et al*., 2014; Machado, 2016; Machado *et al*., 2021). In wild populations from Uruguay, only the form *Psidium cattleyanum* f. *lucidum* is present, and cytotypes with 2n = 55, 66, 77, and 88 have been reported (Speroni et al., 2017). The species exhibits diplosporous apomixis, and unreduced embryo sacs frequently contain more than two nuclei in the central cell (Souza-Pérez and Speroni, 2017). FCSS of seeds derived from controlled crosses in cultivated plants revealed approximately 97% apomixis in heptaploid *P. c.* f. *cattleyanum* (red-fruited) maternal plants and around 80% apomixis in octoploid *P. c.* f. *lucidum* (yellow-fruited) maternal plants (Da Luz-Graña et al., 2025). These studies also reported a high frequency of unreduced female gametes and reduced male gametes, whereas unreduced male gametes were detected at a low frequency (3.2%). Pollination is required for fruit development, indicating a pseudogamous apomictic system (Fidalgo and Kleinert, 2009; Souza-Pérez and Speroni, 2017; Speroni *et al*., 2018; Da Luz-Graña *et al*., 2025). However, pollen viability is highly variable (Raseira and Raseira, 1996; Singhal and Kumar, 2008; Hister and Tedesco, 2016; Vishwakarma *et al*., 2020; Souza-Pérez *et al*., 2021). Microsporogenesis is generally regular, though certain abnormalities occur, leading to the production of polymorphic pollen grains, including unreduced and aneuploid types (Souza-Pérez *et al*., 2021).

The current knowledge of the reproductive biology of *Psidium cattleyanum* is primarily based on studies of cultivated plants (Raseira and Raseira 1996; Singhal and Kumar, 2008; Fidalgo and Kleinert, 2009; Hister and Tedesco, 2016; Souza-Pérez and Speroni, 2017; Vishwakarma *et al*., 2020; Souza-Pérez *et al*., 2021; Da Luz-Graña *et al*., 2025), comparing at most two ploidy levels. However, the reproductive mode of *P. cattleyanum* in natural populations remains unknown, particularly regarding its variation across the different ploidy levels observed in the species. This study aims to investigate the reproductive behavior of *Psidium cattleyanum* f. *lucidum* in wild populations from Uruguay exhibiting different ploidy levels. The research questions to be addressed are: 1) What is the predominant mode of reproduction in wild populations of *P. c.* f. *lucidum*? 2) Are there differences in the dominant mode of reproduction according to the ploidy level of the plants? 3) Is the dominant mode of reproduction associated with the patterns of genetic diversity and population structure in Uruguayan populations?

## Materials and methods

### Wild population sampling

Wild populations of *Psidium cattleyanum* were selected to represent the four ploidy levels reported in Uruguay for the yellow fruit form *P. c.* f. *lucidum* (Speroni *et al*., 2017). These populations were named using a combination of the abbreviated locality name and the ploidy level of the plants: LT-5x (32°10’38.9’’S; 53°47’12.1’’W), CM-6x (33° 0.618; 54° 13.112W), LC-6x (33°52’02.8’’S; 53°31’18.1’’W), R13-7x(34°03’13.1’’S; 53°58’40.5’’W), and BA-8x (32°25’44.3’’S; 53°50’6.6’’W), with vouchers specimens CPerez 361, CPerez 389B, MBonifacino 6715, CPerez 409, CPerez 389, respectively, deposited at the Bernardo Rosengurtt Herbarium of Facultad de Agronomía (MVFA), Uruguay. In each population, individual plants were sampled from a broad spatial distribution within the forest, with a minimum distance of four meters between individuals to reduce the likelihood of clonal sampling. The number of plants sampled per population varied depending on population size: 23 in LT-5x, 15 in CM-6x and LC-6x, 16 in R13-7x and 12 in BA-8x. From each selected mother plant, fresh leaves and mature fruits were collected. Fresh leaves were used for both DNA extraction for molecular marker analysis and ploidy determination via flow cytometry. The fruits were processed to isolate and extract seeds for further analysis. Half of the total number of seeds from each fruit were germinated according to the protocol of Bernaschina and Pereyra (2014) and subsequently analyzed for ploidy level and genetic diversity. The other half of the seeds were used to analyze the DNA content of the embryo and endosperm by flow cytometry in all populations to infer their origin and the reproductive pathway involved. Additionally, seeds from a 2n=4x population from Brazil (Ilha do Cardoso-Cananéia-SP, lat−25.066, long−47.906, Voucher: R.M. Machado 15) were used as a control, as this represents the lowest ploidy level reported for the species, and it has been suggested to reproduce sexually (Machado *et al*., 2022).

### Plant ploidy analysis

The ploidy level of the leaves from all mother plants sampled in each population, as well as their respective seedling progeny, was determined using flow cytometry. In each population, between 45 and 60 germinated seedlings were analyzed, depending on sample availability. The number of seedlings was restricted by subsequent DNA analysis, since in many cases the entire seedling was used for extraction. In the case of LC-6x, progeny ploidy was not studied due to the limited development of the seedlings at the time of the study. For estimation of the nuclear DNA content (2C values), nuclei were isolated following the procedure described by Galbraith *et al*. (1983) using the Woody Plant Buffer (Loureiro *et al*., 2007). Nuclei were stained with propidium iodide (50 µg/mL final concentration) and analyzed using a Partec CyFlow® Space flow cytometer (Sysmex-Partec GmbH, Münster, Germany) with data processing in the FloMax® program (Version 2.4d; Sysmex-Partec GmbH, Münster, Germany). Tomato, *Solanum lycopersicum* Lehm. ‘Stupické Rané’ (2C = 1.96 pg) was used as the internal standard. The estimation of nuclear DNA contents (2C value) was calculated by the equation: sample peak mean/standard peak mean × 2C DNA content of standard (pg). The monoploid genome (1Cx value) of *Psidium cattleyanum,* estimated at 0.52 pg was previously determined in our laboratory (Vázquez Medina, 2014).

### Seed embryo:endosperm balance analysis

The seeds were processed individually, mechanically separating the seed coat from the soft tissues containing the embryo and endosperm (Da Luz-Graña *et al*., 2025). The DNA content of each seed was analyzed following the same flow cytometry protocol described above for the leaves. At the start of each flow cytometry session, FL1 gain was adjusted using a progeny leaf sample of the same ploidy level as the maternal plant, with tomato serving as the internal standard. This enabled us to determine the ploidy level of each embryo and assess its variation relative to the maternal plant. Between 150 and 170 seeds were processed, representing five to six mother plants per population, with six to seven seeds sampled from five fruits per mother. For the 4x Brazilian population, 70 seeds were processed from 7 mother plants, with 10 seeds per mother. A minimum of 3.000 nuclei counts were taken from the peak corresponding to the embryo, and 150 nuclei were analyzed from the endosperm. Coefficients of variation (CV) between 3 and 5 were considered acceptable; in exceptional cases, a CV up to 8 was accepted (with very clear peaks and stable fluorescence over time). The ploidy of the embryos was estimated by calculating the number of genomes, based on the average fluorescence value (Flc embryo) in the sample (including only values +/- 20 of the standard maternal ploidy value). These calculations were used to estimate values for the embryo and the endosperm (Fig. 2a).

**Figure 2:**
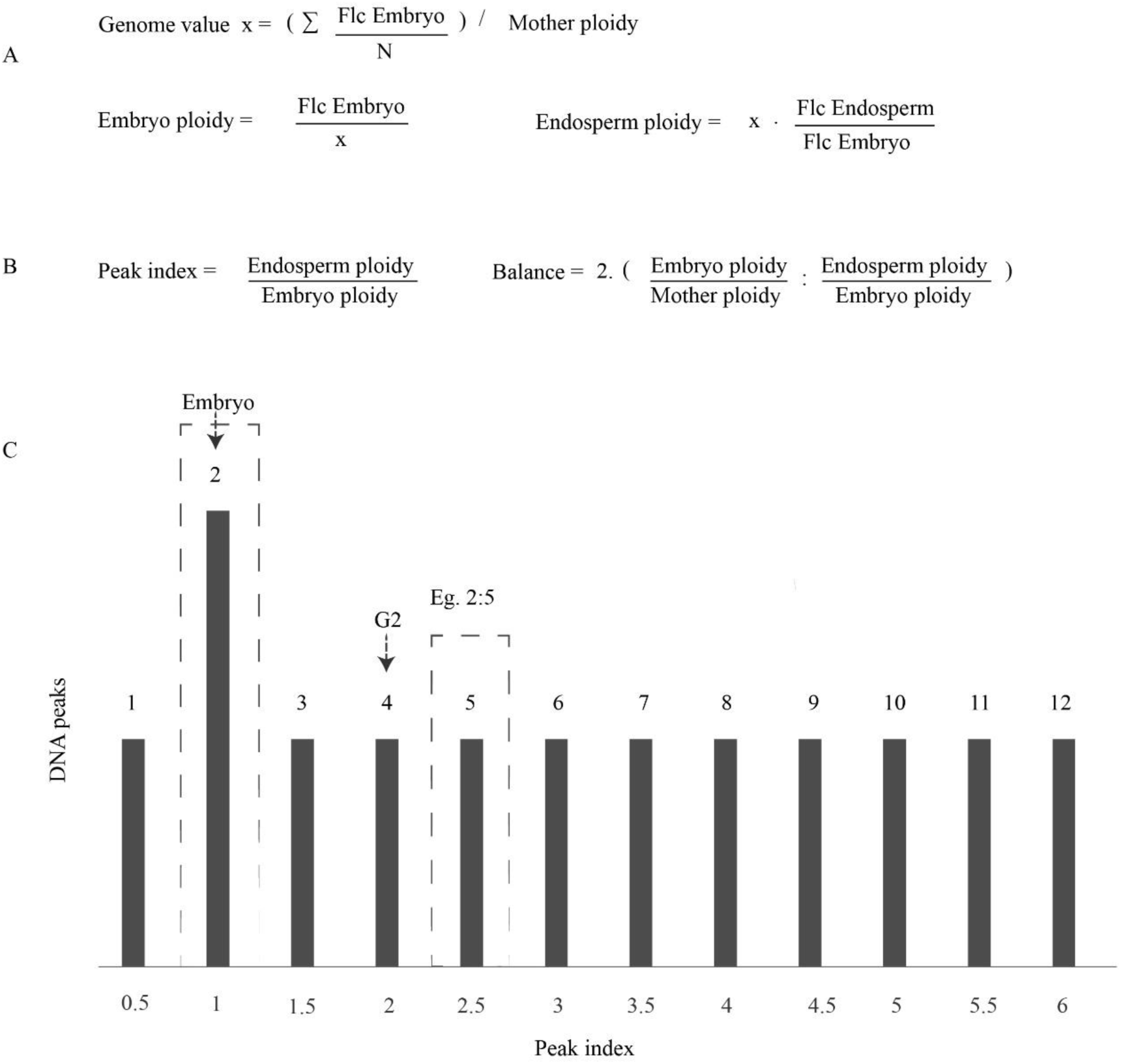
Estimation of the DNA content balance between embryo and endosperm by flow cytometry in seeds (FCSS) of *Psidium cattleyanum* to determine the reproductive origin of both. The analysis is based on the relationship between the fluorescence peaks emitted by the embryo and the endosperm in a histogram. A) Equations used to calculate the ploidy of the embryo and endosperm. B) Peak index and embryo:endosperm balance calculation for each seed. C) Examples of possible peaks in the histogram. The highest fluorescence peak in the histogram corresponds to the embryo (which has the higher number of nuclei). The 2:4 ratio also includes embryo nuclei in the G2 phase of the cell cycle. Between the dashed lines, an example is provided of what the visible peaks would be in a seed with a 2:5 balance.

Embryo and endosperm ploidy were analyzed following the Flow Cytometry Seed Screen (FCSS) methodology by Matzk *et al*. (2000), from which the reproductive mode was inferred. The embryo:endosperm balance was calculated based on the ratio between the embryo and the endosperm fluorescence peak (Fig. 2b), known as the peak index. Seeds with a peak index between 1.3 and 1.7 were considered to be of sexual origin, whereas a peak index of 2.4 or higher indicated apomictic origin (Fig. 2c). Other balances may occur depending on whether the participating gametes are reduced or unreduced, as suggested for the species (Souza-Pérez and Speroni, 2017; Souza-Pérez *et al*., 2021; Da Luz-Graña *et al*., 2025), as well as the presence of a third polar nucleus (Souza-Pérez and Speroni, 2017), and the number of sperm cells fertilizing the central cell (Fig. 2c).

### Statistical analysis of reproductive pathways

Embryo:endosperm ratio frequencies in seeds of apomictic origin were estimated and compared across populations and ploidy levels using contingency tables and Pearson’s Chi-square tests. Sexual reproduction rates were compared among populations— collectively, individually, and pairwise—through deviance analysis and post hoc comparisons using Tukey’s test. Additionally, we compared the types of sexually derived progeny based on their ploidy level relative to that of the maternal plant. For this purpose, embryos were classified as 2C (equal ploidy to the mother plant), 3C (1.5× higher), or 4C (2× higher). Contingency tables were constructed with populations as rows and the frequency of each category as columns. A global test of independence was performed using Pearson’s chi-square test and the multivariate G² (MV-G2) test. Corresponding p-values were calculated and adjusted for multiple comparisons using the Bonferroni correction, in order to control for type I error inflation due to multiple testing. All statistical analyses were conducted using R software (R Core Team, 2020) and InfoStat (Di Rienzo *et al*., 2018).

### SSR amplification and analysis

All leaf DNA extractions were carried out using a CTAB-based method (Viruel and Hormaza, 2004). DNA quantity and quality were checked using a Nanodrop ND-1000 UV-visible spectrophotometer, then diluted to 10 ng μl^−1^ with water. PCR amplifications followed the protocol of Viruel and Hormaza (2004) with amplification conditions based on those described by Machado *et al*. (2021) for SSR-specific primers and Tuler *et al*. (2015) for SSR primers cross-amplified from *P. guajava*. Amplifications were performed in a thermocycler (Bio-Rad Laboratories, Hercules, CA, USA). Forward primers were labeled with a fluorescent dye at the 5’ end. The PCR products were analyzed using capillary electrophoresis with a Beckman Coulter Genome LabTM GEXP capillary DNA analysis system for genotyping. PCR amplicons were prepared and denatured at 90°C for 120 s, injected at 2.0 kV for 30 s, and separated at 6.0 kV for 35 min during capillary electrophoresis. Each PCR and capillary electrophoresis was repeated at least twice to ensure the reproducibility of the results.

A preliminary screening was made with leaves from all mother plants in each population using microsatellite (SSR) markers transferred from *Psidium guajava* (Risterucci *et al*., 2005; Guavamap, 2008) and specific markers for *Psidium cattleyanum* (Machado *et al*., 2021) [Table 1S, **supplementary information**]. The six most informative markers (three from each source) were selected, based on the presence of polymorphisms and the clarity of the peaks (Table 1). Progeny plants were analyzed using the selected SSR markers, with 140-160 seedlings in each population.

**Table 1:**
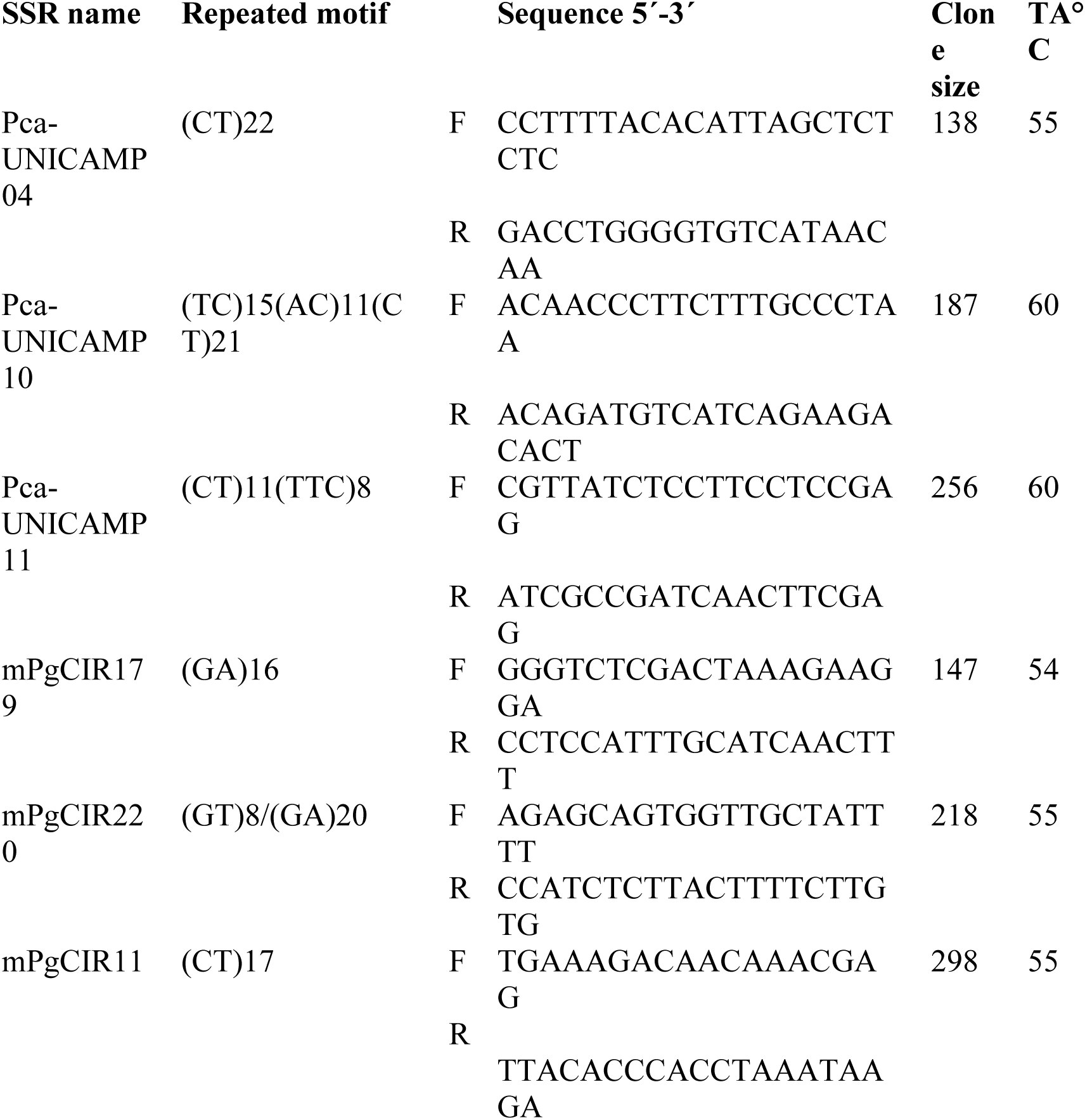
SSR primers used for the detection of polymorphisms in the mother plants and progenies of *Psidium cattleyanum* f. *lucidum*.

### Microsatellite validation and genetic diversity analysis

For primer characterization, microsatellites were treated as dominant markers (presence= 1 and absence= 0). For each marker, the number of bands (NB), polymorphism content (PIC; Botstein *et al*., 1980), and discriminatory power (DP) were calculated, as well as the number of private bands (PB) per population. Population genetic parameters were estimated by calculating Shannon, Simpson and Nei indices for each population (mother plants with their progeny) using the *poppr* and *polysat* packages on the R 3.4.0 platform. Genetic distance among individuals was computed using the Bruvo distance method (Bruvo *et al*., 2004).

To infer the population structure of the individuals studied, we used a Discriminant Analysis of Principal Components (DAPC; Jombart *et al*., 2010), which uses a nonparametric approach, free from Hardy-Weinberg constraints, performed in the *adegenet* package (Jombart, 2008). Two approaches were used: (1) the first DAPC analysis was performed by providing information for five groups (populations of *P. cattleyanum*); and (2) the number of clusters was evaluated using the *find.clusters* function. The maximum number of clusters assumed was 15. The optimal number of clusters was estimated using the Bayesian information criterion (BIC), and both DAPC results were presented as dot plots.

A molecular variance analysis (AMOVA) was performed using the *poppr* package (Kamvar *et al*., 2014), to investigate the partitioning of genetic variance within and between populations, as well as among pre-defined groups based on ploidy level and reproductive system (considering the most frequent pathways observed in the population according to the embryo/endosperm balance analysis). Prior to this analysis, the *clone.correct* function was applied to our matrix to remove potential biases from cloned genotypes.

## Results

### Ploidy level of mother plants, seedlings and embryos

All plants studied in each population exhibited the same ploidy level: LT-5x, pentaploid; CM-6x and LC-6x, hexaploid; R13-7x, heptaploid; and BA-8x, octoploid. Among the progeny, 70 to 90% retained the same ploidy level as their respective mother plants. The percentage of seedlings with different ploidy levels varied depending on the population (Fig. 3a), and in all cases, it was higher than that observed in the mother plant. Populations with odd ploidy levels presented the highest number of new cytotypes in their progeny. In the pentaploid population, three additional cytotypes (2C=6x, 7x, 8x) were detected, while in the heptaploid population, two new cytotypes (2C=10x, 11x) were observed (Fig. 3a). In contrast, populations with even ploidy levels showed only one additional cytotype, different from the maternal one, the two hexaploid populations produced also seedlings with 2C= 9x and the octoploid population included 2C= 12x individuals (Fig. 3a). Flow cytometry analysis of embryo DNA content revealed several cytotypes not observed among the progeny seedlings (Fig. 3b). This analysis also detected ploidies lower than those of the maternal plants (LT-5x 2C= 3x, LC-6x 2C = 3x; R13-7x 2C= 6x; BA-8x 2C= 4x) as well as new higher ploidy levels (LT-5x 2C= 10x, LC-6x 2C= 9x; R13-7x 2C= 14x). Populations with odd ploidies, LT-5x, and R13-7x, exhibited the greatest diversity of ploidy levels (Fig. 3a and b). Overall, considering both seedlings and embryos, progeny ploidy levels across all populations ranged from 3x to 14x.

**Figure 3.**
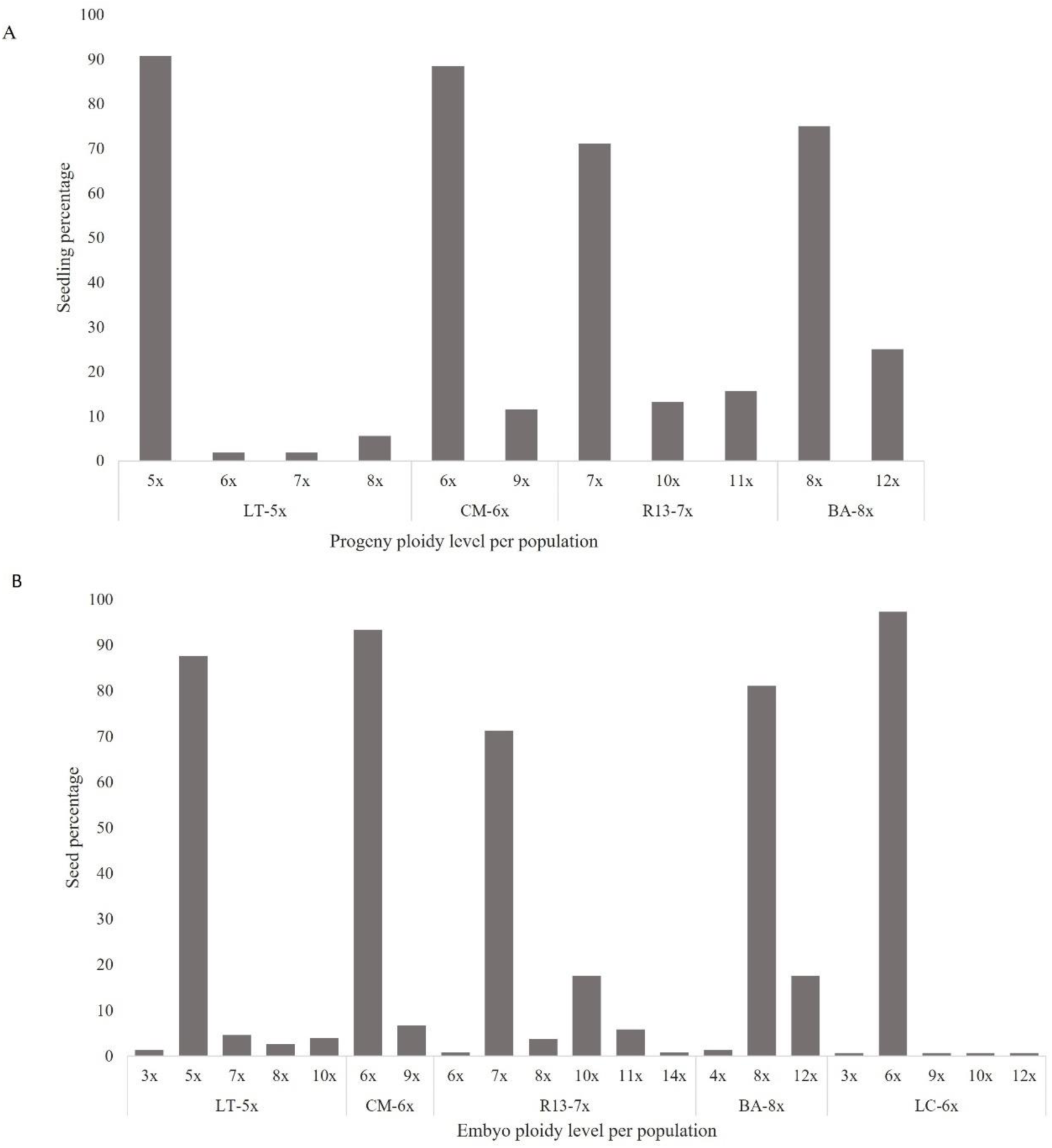
Ploidy level of progeny seedlings and embryos in *Psidium cattleyanum* f. *lucidum* wild populations. A) Percentages of ploidy levels found in progeny seedlings from four populations (LT-5x, CM-6x, R13-7x, and BA-8x). B) Percentages of ploidy in embryos in seeds from five populations (LT-5x, CM-6x, R13-7x, BA-8x, and LC-6x). In the populations, *x* represents the basic chromosome number.

### Embryo: endosperm DNA content balance

A total of 810 seeds were analyzed for embryo:endosperm DNA content. Of all the processed seeds, approximately 5% were empty, and 1% did not show the endosperm peak in the flow cytometry histograms. In most of the analyzed seeds, the histograms displayed three distinct peaks of emitted fluorescence: the embryo peak, corresponding to the population of nuclei in G0/G1, which was identified as the highest peak due to its greater number of nuclei; the double embryo peak corresponding to the population of nuclei in the G2 phase of the cell cycle; and the peak corresponding to the nuclei of the endosperm (Fig. 4). However, in several seeds, two endosperm peaks were observed (Fig. 4b, d, h), and occasionally, additional distant peaks were detected, likely corresponding to endoduplications events of the embryo or endosperm. The analysis of histograms from individual seeds revealed a variety of endosperm and embryo origins in each population, determined by the embryo:endosperm balance they exhibited (Fig. 5). The most frequent balances in the populations were 2:5, 2:5:6, 2:6, 2:3:5 (Fig. 4 a, b, c, d respectively), corresponding to the apomictic pathway (Fig. 5), and 2:3, 3:5, 3:5:2 (Fig. 4 f, g, h respectively), corresponding to the sexual pathway (Fig. 5). Some other balances were present at very low frequencies, in both the apomictic and sexual pathways (Fig. 5). Populations exhibited significantly different distributions of embryo:endosperm balance types, with proportions depending both on ploidy level and population identity (p-values <0.0001).

**Figure 4:**
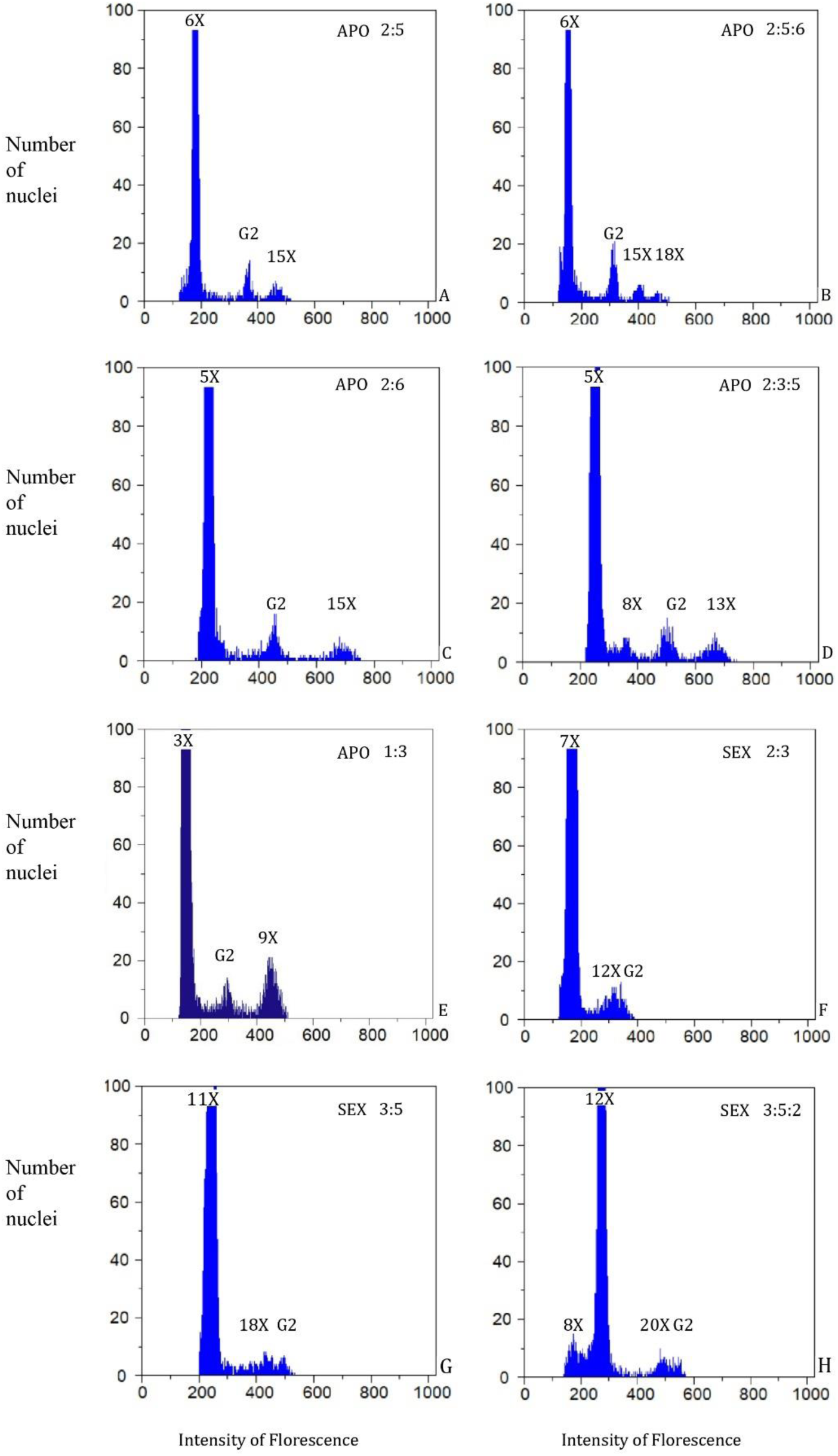
Representative histograms (*x-*axis: Intensity of fluorescence; *y-*axis: number of nuclei) from flow cytometry seed screen (FCSS) analysis from *Psidium cattleyanum* f. *lucidum* wild populations. The different balances between the DNA contents of the embryo and endosperm nuclei correspond to the peaks, presented as embryo: endosperm1: endosperm2 (if it is the case), G2 peak was not included. A) 2:5 balance, population CM-6x; B) 2:5:6 balance, population CM-6x; C) 2:6 balance, population LT-5x BA-8x; D) 2:3:5 balance, population LT5x; E) 1:3 balance, population LT-5x; F) 2:3 balance, population R13-7x; G) 3:5 balance, population R13-7x; H) 3:5:2 balance, population. APO: Apomixis. SEX: Sexuality. G2: embryo nuclei in the G2 phase of the cell cycle.

**Figure 5:**
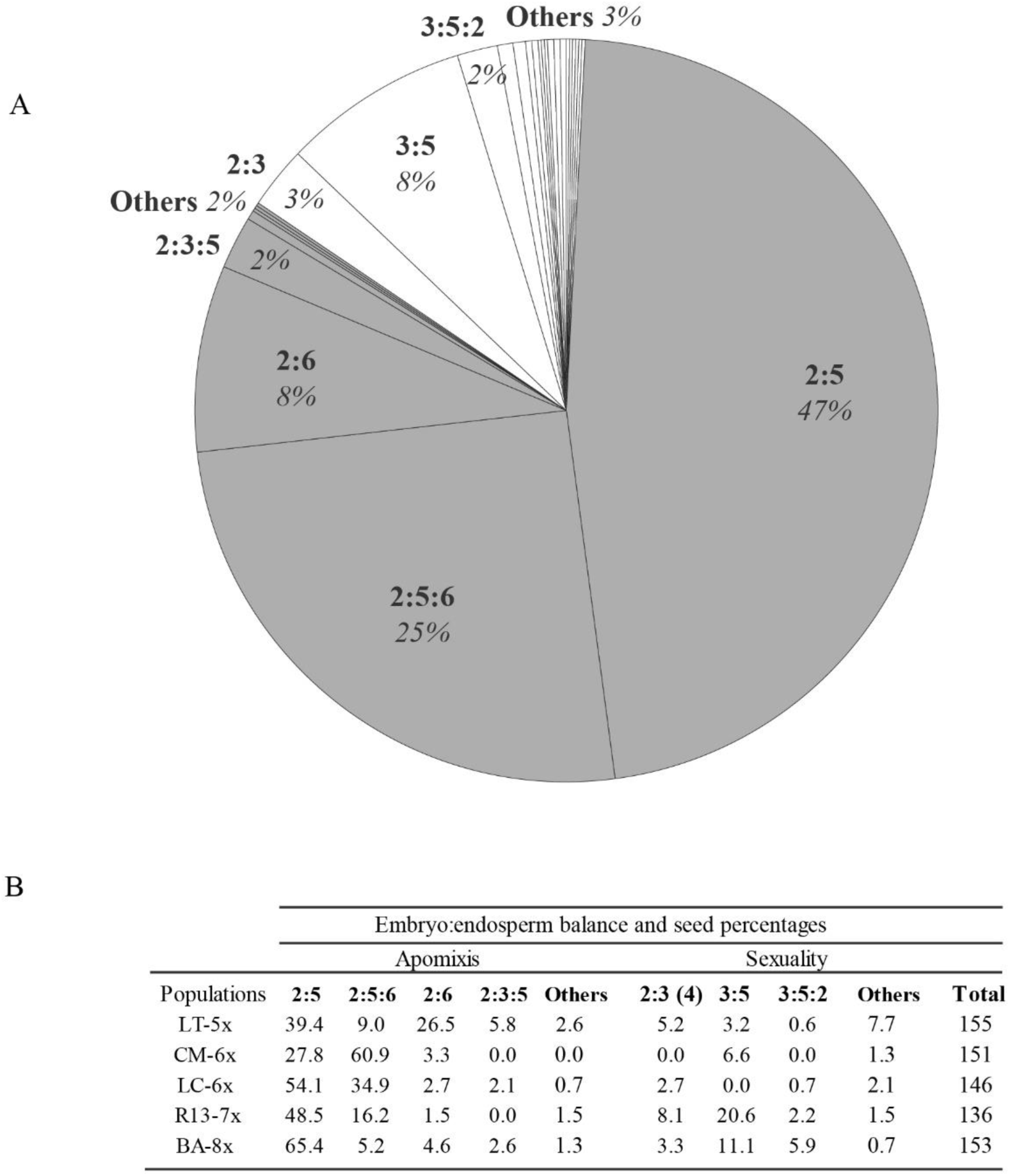
Embryo:endosperm balances found in seed analysis of *Psidium cattleyanum f. lucidum* wild populations from Uruguay. A) Relative frequency of the balances found in five populations, only those percentages greater than 1% were identified. The balances are presented as embryo: endosperm1: endosperm2 (if it is the case). In gray are the ones resulting from apomixis, and in white, those resulting from sexual reproduction. B) Absolute frequency of these balances in each population.

### Reproductive pathways in wild populations

In the analyzed wild populations of *Psidium cattleyanum* f. *lucidum*, apomixis was the most frequent reproductive pathway, ranging from approximately 70 to 95% of the seeds analyzed (Fig. 6). The highest frequency of apomixis was observed in the hexaploid populations (LC-6x and CM-6x). Although apomixis was dominant across all populations, sexual reproduction was detected in each population, with the highest frequencies found in those populations with the highest ploidy levels: R13-7x (32%) and BA-8x (21%). In the Brazilian population used as control (Icar-4x), sexual reproduction was the dominant pathway, occurring in 99% of cases. Comparative Tukey tests based on ploidy and population within the Uruguayan populations revealed significant differences in sexual reproduction rates among some populations (Fig. 6), notably between R13-7x and BA-8x compared to LT-5x, CM-6x, and LC-6x. When comparing by ploidy levels, significant differences were observed between 5x and 6x populations [Table 2S, **supplementary information**].

**Figure 6.**
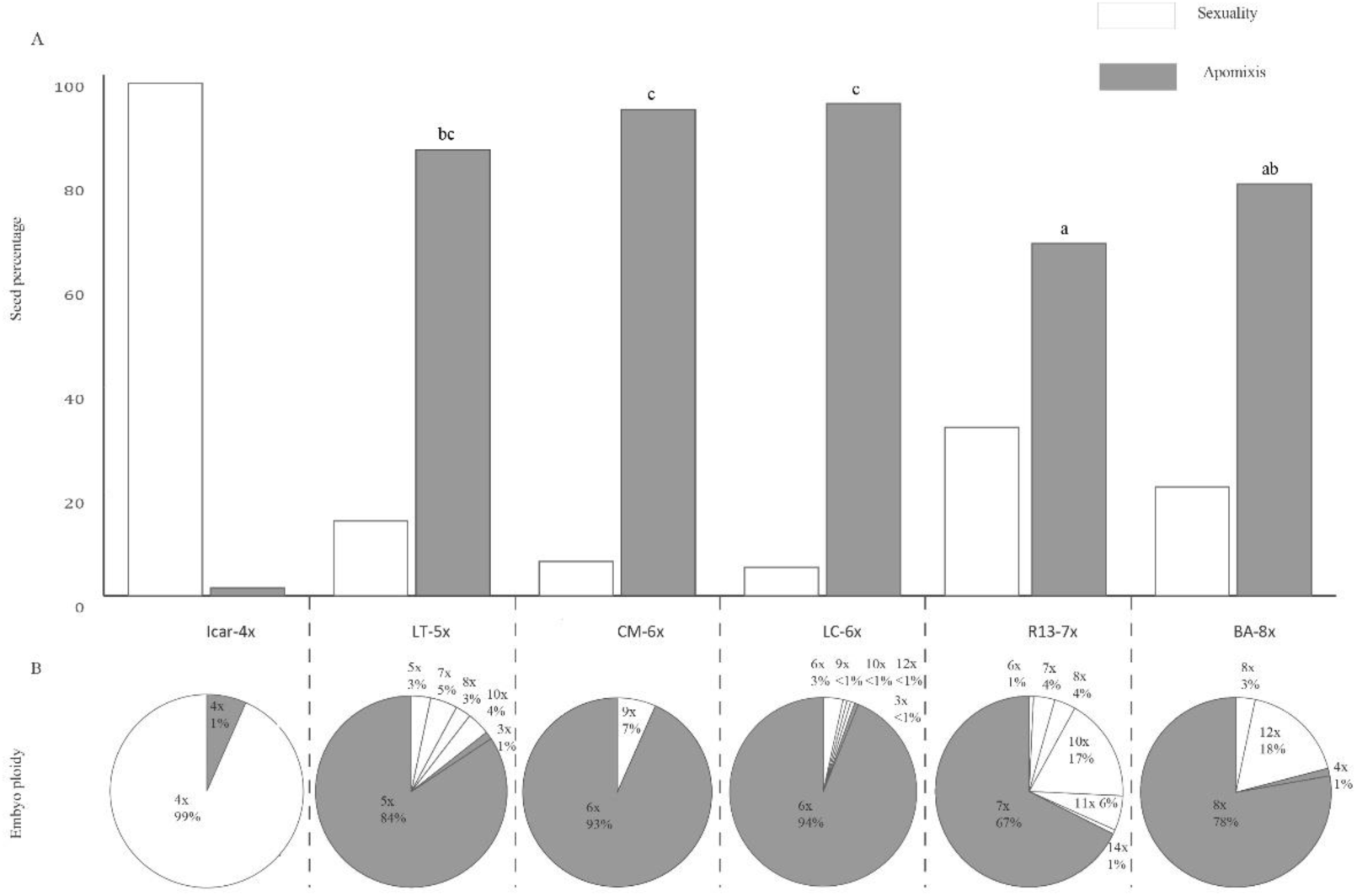
Frequency of reproductive modes and embryo ploidy of *P. cattleyanum* f. *lucidum* wild populations. A) Percentages of apomictic (gray) and sexual (white) seeds in populations with different ploidy levels (Icar-4x, LT-5x, CM-6x, R13-7x, and BA-8x). B) Percentages of embryos and their ploidy level in each population, of apomictic (gray) and sexual (white) origin. Different letters indicate significant differences according to Tukey’s test.

Sexually derived embryos exhibited ploidy levels equal (2C) or higher (3C, 4C) than those of the mother plants (Fig. 6b). Type 2C embryos, which maintain the mother plant ploidy, originating from the fusion of reduced gametes, were observed in 2-20% (LT-5x, CM-6x, R13-7x and BA-8x) and 40% (LC-6x) of sexually originated seeds. Most seeds originated from unreduced embryo sacs, fertilized either by a reduced sperm cell (producing 3C embryos) or, less frequently, by unreduced sperm cells (resulting in 4C embryos). Sexual embryos of type 3C were the predominant type in all populations: LT-5x=7x, 8x; CM-6x=9x; LC-6x=9x, 10x; R13-7x=8x, 10x, 11x; BA-8x=12x (Fig. 6b). The populations that presented 4C embryos were LT-5x, LC-6x, and R13-7x. The frequency distribution of embryo types (2C, 3C, 4C) differed significantly among populations (p<0.0083), with LT-5x showing notable differences compared to R13-7x and BA-8x [Table 3, **supplementary information**].

### SSR marker characterization

All six selected SSR loci proved to be informative for the studied populations. Their discriminatory power (their ability to differentiate between individuals or genotypes within a population) varied between 0.84 to 0.87. The PIC values obtained for the six markers ranged from 0.81 to 0.90, indicating a high level of informativeness. For the six loci analyzed [Table 4, **supplementary information**], the number of amplified bands per locus ranged from 11 to 29. The number of exclusive bands, i.e. observed in only one population, ranged from 6 to 26. The LC-6x population showed the highest number of exclusive bands, with a total of 26.

### Population diversity and structure

A total of 191 SSR bands were amplified from the six microsatellite loci (Table 2). The number of bands per population ranged from 29 to 50. The genetic diversity parameters, including the Shannon, Simpson, and Nei indices, estimated for each population ranged from 2.43 to 3.34, from 0.779 to 0.931, and from 0.0779 to 0.0971, respectively (Table 2). Genetic diversity analyses revealed high genetic diversity in the LC-6x population (Shannon index = 3.34, Simpson index = 0.931, Nei index = 0.0790) which was the population with the most observed polymorphisms, especially with markers transferred from *P. guajava*. The BA-8x population presented the highest values of genetic variability (Nei index = 0.0971), although it was the population with the lowest number of polymorphic individuals. The population with the lowest diversity indices was R13-7x, with very few polymorphisms and only in one locus. All five populations of *P. cattleyanum* f. *lucidum* exhibited a low quantity of genotypes (Global: 213 of 786 individuals).

**Table 2.** Parameters of genetic diversity estimates for the Uruguayan populations of *Psidium cattleyanum* evaluated in this study. N number of individuals; MLG multilocus genotypes number; NB number of bands.

| Population | N | MLG | Shannon Index | Simpson Index | Nei index | NB |
| --- | --- | --- | --- | --- | --- | --- |
| LT-5x | 163 | 42 | 2.64 | 0.838 | 0.0891 | 34 |
| BA-8x | 162 | 38 | 2.45 | 0.779 | 0.0971 | 41 |
| CM-6x | 159 | 37 | 2.59 | 0.837 | 0.0806 | 37 |
| R13-7x | 162 | 39 | 2.43 | 0.782 | 0.0779 | 29 |
| LC-6x | 140 | 57 | 3.34 | 0.931 | 0.0790 | 50 |
| Global | 786 | 213 | 4.28 | 0.965 | 0.2183 | 191 |

The DAPC analyses are presented in Fig. 7. Delimitation of populations was clear in both tested approaches, indicating that the five sampled populations are genetically distinct from each other. A heatmap based on Bruvo’s distance clustered the individuals together into populations, although some individuals were mixed [Fig. 1S, **supplementary information**].

**Figure 7.**
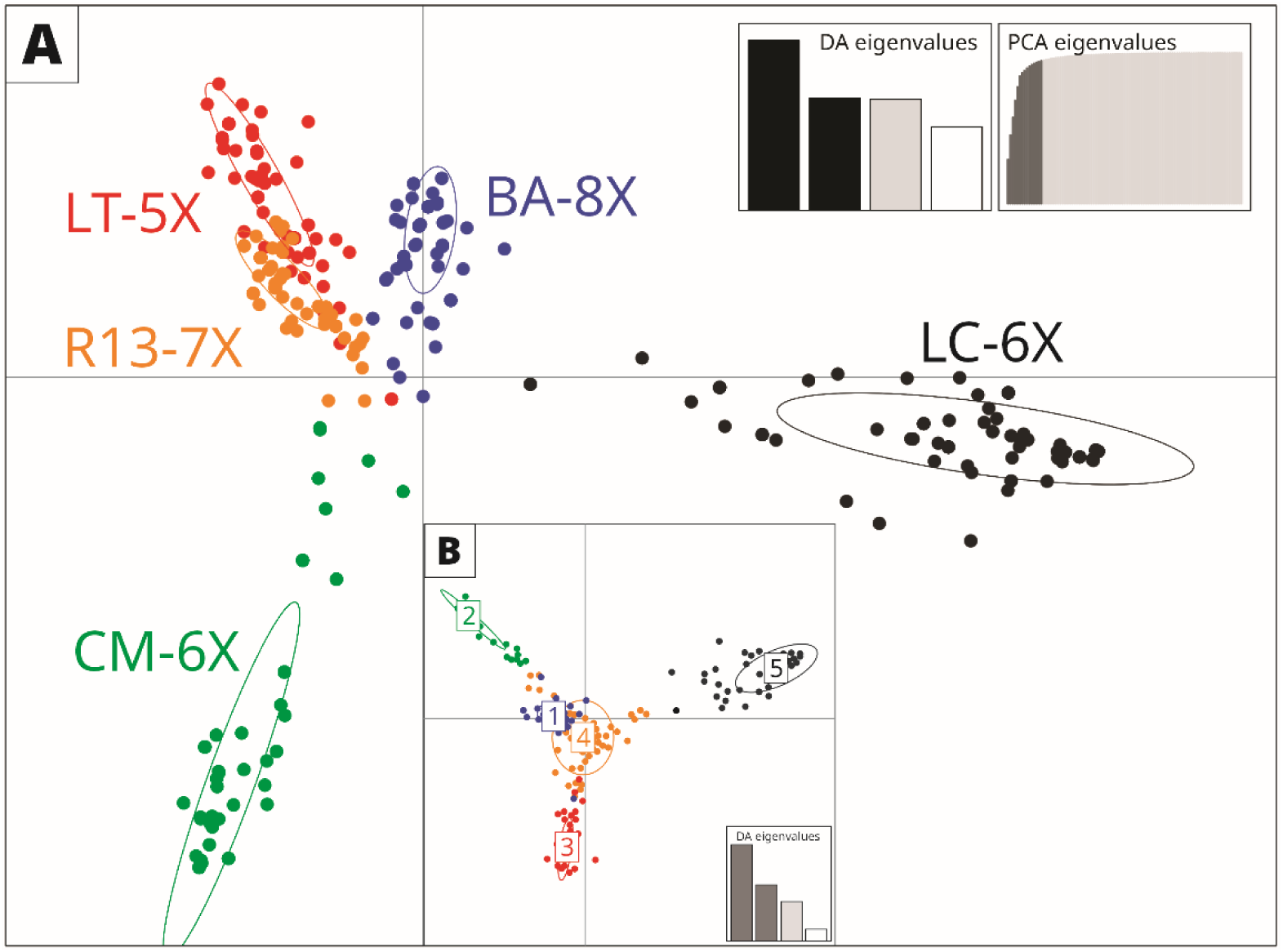
Population structure of *Psidium cattleyanum* individuals inferred by Discriminant Analysis of Principal Components DAPC (A) Analysis was performed by providing information for five groups (populations of *P. cattleyanum*); (B) The number of clusters was evaluated using the *find.clusters* function. Colors identified individuals from each population.

The AMOVA showed that a large fraction of genetic variation was partitioned among populations grouped by types of reproduction (ϕST = 0.573 and 0.581, p < 0.001; Table 3), suggesting that genetic differentiation was related to the mode of reproduction. The analysis also showed significant genetic structure between reproductive types (ϕCT = 0.585, p < 0.001; Table 3)

**Table 3.**
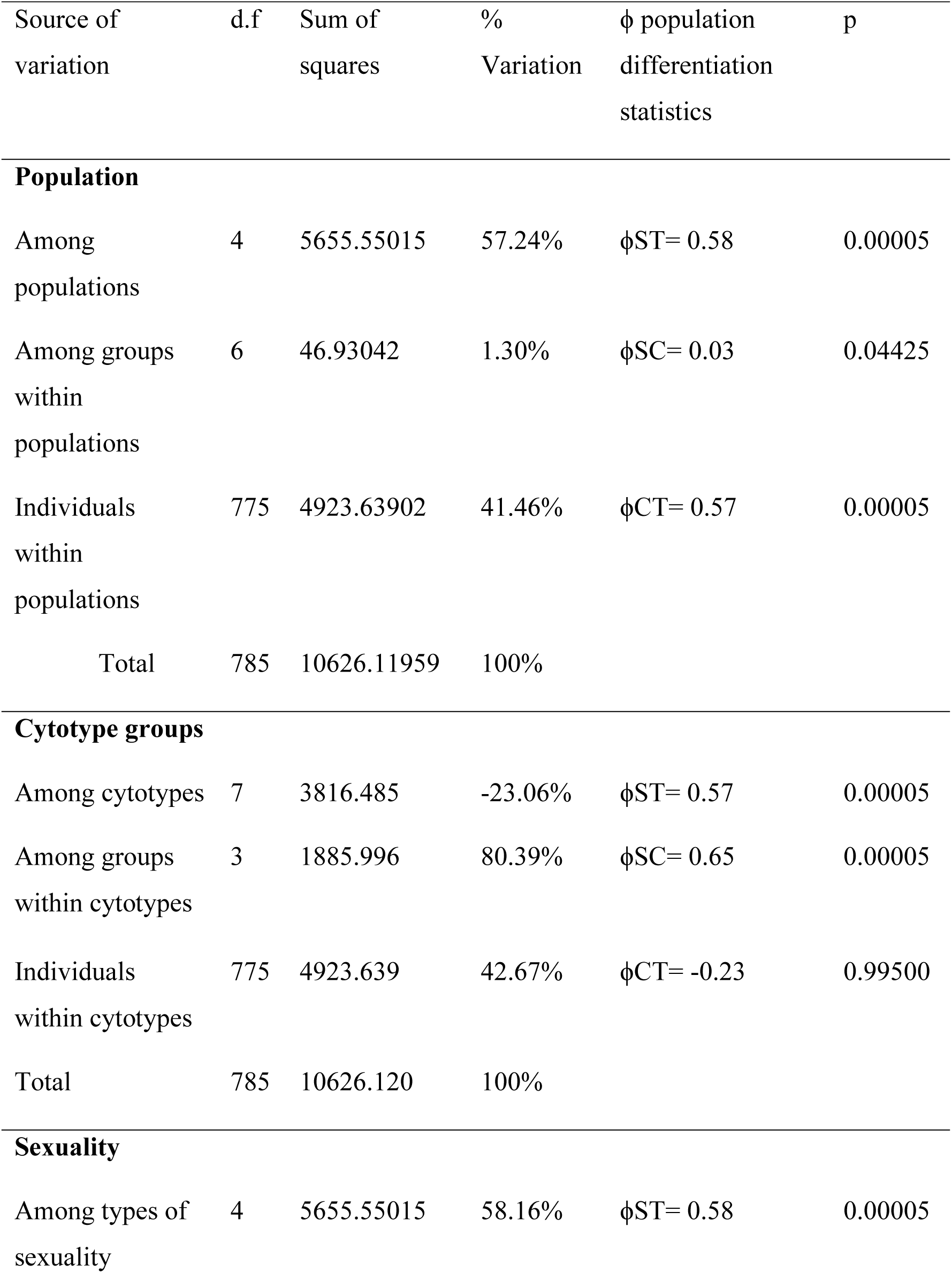
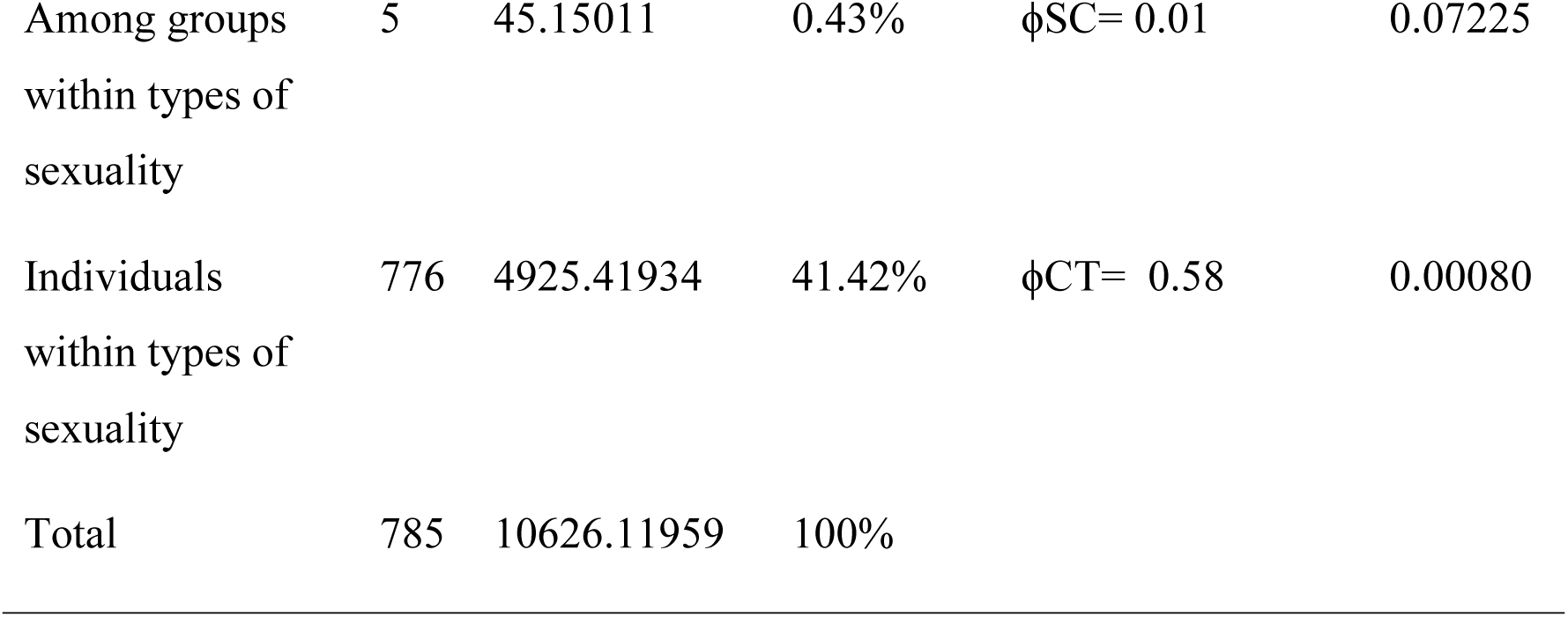
Analysis of molecular variance (AMOVA) based on six microsatellite markers in *Psidium cattleyanum*. Three tests were performed: (1) the population being higher hierarchy, (2) cytotype groups being higher hierarchy and (3) types of reproduction being higher hierarchy.

## Discussion

In this study, we investigated the mode of reproduction in wild populations of *Psidium cattleyanum* f. *lucidum,* including the four ploidy levels present in Uruguay (2n = 5x, 6x, 7x, 8x). Two complementary approaches were used: seed scanning by flow cytometry (FCSS) and assessment of progeny genetic variability using microsatellite markers (SSRs). We found that pseudogamous apomixis is the predominant mode of reproduction across the studied populations. Significant differences were observed among these populations in the frequency of reproductive modes (apomictic vs. sexual) as well as in the proportions of progeny types (embryo ploidy levels) and seed types (embryo:endosperm DNA content balance). The genetic variability found in the progeny indicates that clonal reproduction is predominant in seeds of Uruguayan *P. c.* f. *lucidum* populations. The integration of cytogenetic and molecular data reveals that the mode of reproduction seems to be the driving force shaping the genetic structure. This study highlights the complexity of the species’ reproductive system, which is influenced by population-specific ploidy levels and reproductive strategies.

### Reproductive modes in Psidium cattleyanum f. lucidum

Both the FCSS and SSR approaches show that the majority of seed progeny is clonal, with apomixis being the dominant mode of reproduction in all populations. Sexual reproduction is present in 6–33% of individuals within populations, supporting the presence of facultative apomixis previously suggested for the species through various approaches, such as hand-pollinated crosses (Raseira and Raseira, 1996; Da Luz-Graña *et al*., 2025) and embryological studies (Souza-Pérez and Speroni, 2017) in cultivated plants, as well as genetic diversity analyses in natural populations from Brazil (Machado *et al*., 2021, 2022). These findings reinforce the idea that facultative apomixis is the prevailing condition among apomictic angiosperms (Krahulcová *et al*., 2004; Tucker and Koltunow, 2009). The presence of at least some sexual reproduction in apomictic species is essential to prevent the accumulation of deleterious mutations that could otherwise lead to the extinction of strictly apomictic lineages (Hojsgaard and Hörandl, 2015; Hodač *et al*., 2019; Hörandl, 2024). Therefore, the coexistence of both systems provides an evolutionary advantage, combining the benefits of maintaining well-adapted clonal genotypes via apomixis, while also allowing occasional recombination and genetic variation to respond to environmental changes (Hörandl, 2024).

The principal pathways of embryo formation in gametophytic apomixis are present in nearly all studied populations of *Psidium cattleyanum* f. *lucidum*: diploid parthenogenesis, haploid parthenogenesis, sexual reproduction via reduced gametes, and sexual reproduction via unreduced gametes (Nogler, 1984). A cytological interpretation of the various reproductive events identified (based on embryo:endosperm balance) is presented in Figure 8a for apomictic seeds and in Figure 8b for sexually derived seeds. These interpretations consider both reduced and unreduced embryo sacs and pollen grains in both reproductive pathways. We based our interpretations on common features observed in angiosperms and specifically reported in this species: (1) two polar nuclei that jointly participate in endosperm formation, fusing either before or at the time of fertilization (Nogler, 1984; Dobeš *et al*., 2013b; Souza-Pérez and Speroni, 2017; Da Luz-Graña *et al*., 2025) ; (2) the central cell may be fertilized by one or two sperm nuclei (Talent and Dickinson, 2007; Qu *et al*., 2010; Galla *et al*., 2011; Da Luz-Graña *et al*., 2025); and (3) the occurrence of a third polar nucleus in the embryo sac (Nogler, 1984; Talent and Dickinson, 2007; Souza-Pérez and Speroni, 2017; Da Luz-Graña *et al*., 2025).

**Figure 8:**
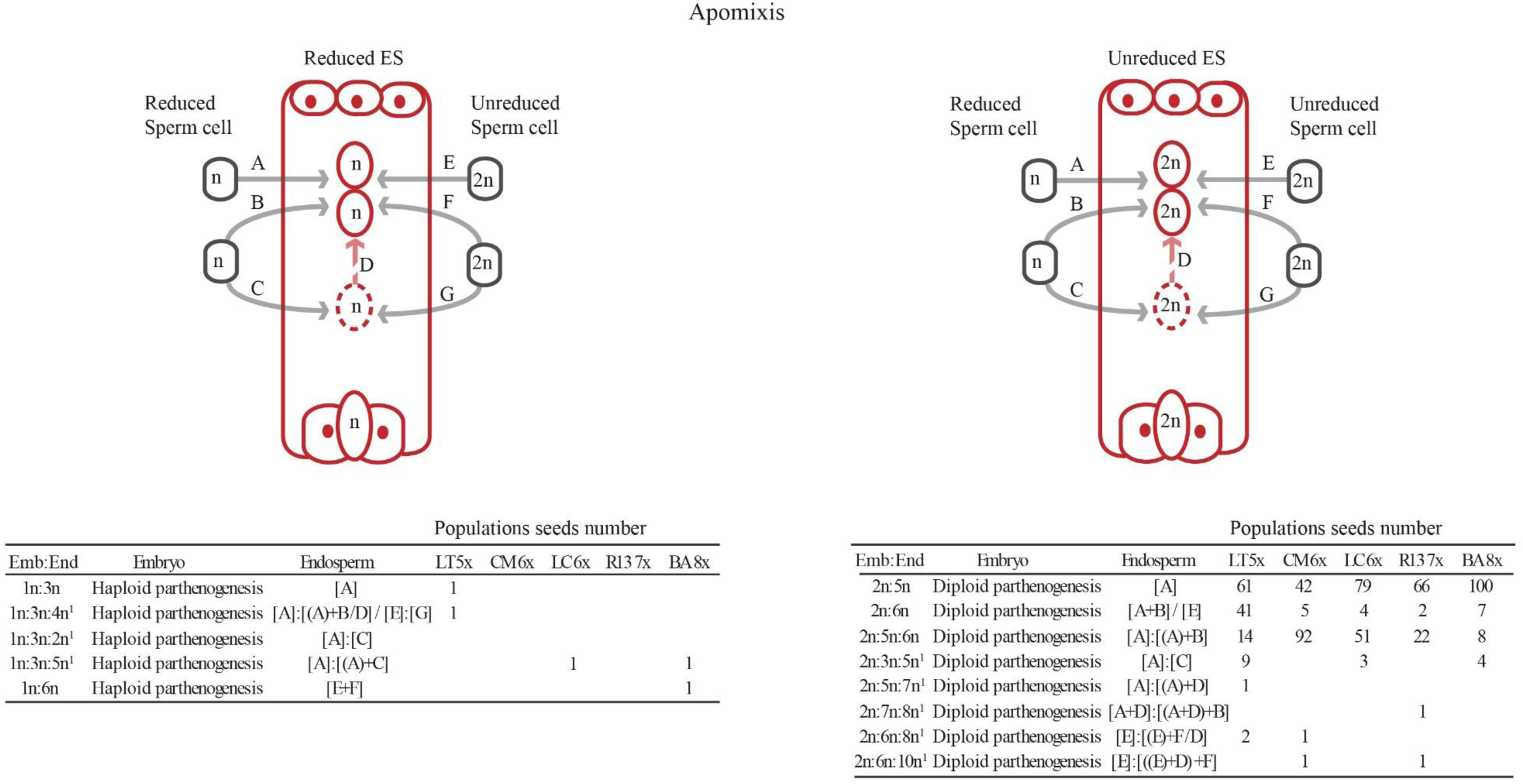
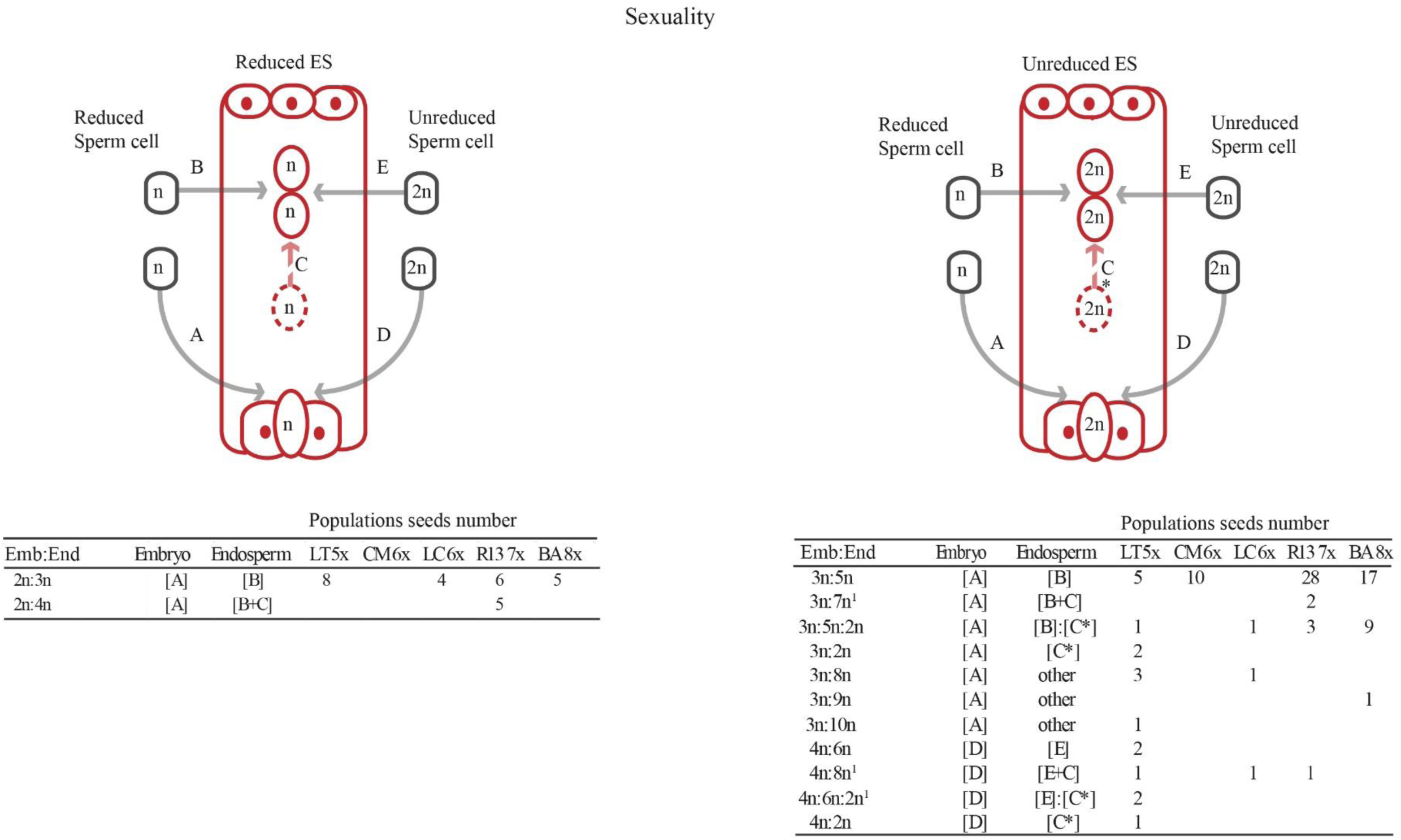
Embryo:endosperm balances found in seeds of *Psidium cattleyanum* f. *lucidum* and representation of seed origin, based on the type of embryo sac and reduced (n) or non-reduced (2n) sperm cells involved. Uppercase letters identify cell fusion (fertilization or not); “[]” encompasses all steps occurring for the formation of the different ploidies in the balances (peaks in the histogram); “()” identifies the formation of preliminary tissue; “+” denotes tissue fusion; “:” separates peaks observed in the histogram; “1” indicates the involvement of a third polar nucleus. It is assumed that two polar nuclei fuse, and the third may join the set, fuse later, or not participate. **A) Cases of the apomictic reproductive pathway.** The reduced embryo sac is shown on the left, and the non-reduced embryo sac is on the right. All the balances observed are represented, along with the number of cases for each population, classified by the type of reproductive pathway, embryo sac type, and the sperm cell that originated it. **B) Cases of the sexual reproductive pathway.** The reduced embryo sac is shown on the left, and the non-reduced embryo sac is on the right. All the balances observed are represented, along with the number of cases for each population, classified by the type of reproductive pathway, embryo sac type, and the sperm cell that originated it.

Several of the embryo sacs (ES) and pollen grains (PG) characteristics identified in the present study are consistent with those reported by Da Luz-Graña *et al*. (2025) in seeds derived from hand-pollinated crosses between cultivated plants of *Psidium cattleyanum* f. *cattleyanum* (7x) and *P. c.* f. *lucidum* (8x). In the wild populations, embryo sacs (ES) were unreduced in more than 90% of cases (Table 4). Reduced embryo sacs were observed in some sexually originated embryos and in haploid parthenogenesis. A third nucleus in the central cell was evidenced in unreduced ES and, occasionally, in the reduced ones. Pollen grains were predominantly reduced (Table 4, Figures 8a and b), in accordance with the high frequency of meiosis observed in cytological studies (Souza-Pérez *et al*., 2021). The ploidy level of pollen donors was considered the same as that of the mother plant, based on the homogeneity of ploidy levels within populations. Although the presence of aneuploid pollen grains is expected in this species (Souza-Pérez *et al*., 2021) and is common in other diplosporous species (Pagliarini, 2000; Singhal and Kumar, 2008; Cornaro *et al*., 2023), in this analysis we only considered reduced and unreduced pollen grains, as the approaches used do not allow clear discrimination between aneuploidy and minor differences in genome size between the embryo and endosperm (Šarhanová *et al*., 2012).

**Table 4.** Percentages of reduced (n) and unreduced (2n) embryo sacs (ES) and pollen grains (PG). Percentage of embryo sacs containing three polar nuclei (3pn)

|  | ES |  | ES |  | PG |  |
| --- | --- | --- | --- | --- | --- | --- |
|  | n | 3np | 2n | 3np | n | 2n |
| LT-5x | 4.6 | 0.7 | 95.4 | 7.8 | 84.3 | 15.7 |
| CM-6x | 0.0 | 0.0 | 100.0 | 1.3 | 95.4 | 4.6 |
| LC-6x | 4.1 | 0.7 | 95.9 | 2.1 | 95.9 | 4.1 |
| R13-7x | 8.1 | 0.0 | 91.9 | 1.5 | 97.8 | 2.2 |
| BA-8x | 4.6 | 0.7 | 95.4 | 2.6 | 99.3 | 0.7 |

### Embryo origin

Embryos are predominantly formed via diploid parthenogenesis from unreduced embryo sacs (ES), consistent with diplosporous apomixis as proposed in previous ontogenetic studies of the species (Souza-Pérez and Speroni, 2017). Haploid parthenogenesis, also referred to as non-recurrent apomixis (Maheshwary, 1950), has been previously observed in the species (Da Luz-Graña *et al*., 2025) and also in several apomictic plant groups (e.g., *Potentilla puberula* (Dobeš *et al*., 2013b), *Rubus* subgen. *Rubus* (Šarhanová *et al*., 2012, 2024); *Crataegus* (Talent and Dickinson, 2007); *Pilosella rubrum* (Krahulcová *et al*., 2004), though always at low frequency. Sexual embryos arise from both reduced and unreduced egg cells, with the latter resulting in the so-called BIII hybrids (embryos with ploidy levels higher than the maternal plant), which were predominant among both types in nearly all populations studied (in this study, progeny with 3C and 4C ploidy levels).

The comparison of sexual-origin embryo types based on their ploidy level (2C, 3C, and 4C) revealed statistically significant differences among LT-5x, R13-7x, and BA-8x populations. The LT-5x and R13-7x populations exhibited a higher number of distinct ploidy levels in the embryos and higher percentages of 4C embryos (Figure 7, Figure 8b). These populations, with odd ploidy levels, may produce gametes of various ploidy levels, or chromosome numbers, due to deviations in meiosis, as previously observed in this species (Singhal *et al*., 1985; Souza-Pérez *et al*., 2021) and other angiosperms (Asker and Jerling, 1992; Rotreklová and Krahulcová, 2016; Farco and Dematteis, 2017). This gamete disparity results in a greater range of ploidy combinations involved in embryo formation.

On the other hand, the R13-7x and BA-8x populations exhibited the highest proportions of 3C progeny (BIII hybrids), concomitant with the highest levels of sexual reproduction. The proportion of BIII hybrids in these populations (heptaploids > 20%; octoploids > 10%) aligns with those reported for octoploid *P. c.* f. *lucidum* plants (16.5%) but contrasts with those documented for heptaploid *P. c.* f. *cattleyanum* plants (0.8%) (Da Luz-Graña *et al*., 2025). Notably, those values are relatively high compared to other apomictic species, which generally range from 1% to 3% (León-Martínez and Vielle-Calzada, 2019). A predominance of BIII hybrids has also been observed in other facultative diplosporic apomixis species such as *Hieracium* s.s (Mráz and Zdvořák, 2019), and in the progeny of facultatively apomictic hexaploid *H. rubrum,* when crossed to tetraploid *H. pilosella* (Krahulcová *et al*., 2004). These studies suggest that unreduced female gametes in polyploids may have a higher fertilization success rate due to meiotic abnormalities (Mráz and Zdvořák, 2019), as well as differences in maternal ability to generate unreduced gametes capable of fertilization (Krahulcová *et al*., 2004). Our study further contributes to this evidence by showing that, when comparing multiple ploidy levels within the same species, unreduced female gametes display a greater fertilization capacity at higher ploidy levels.

### Endosperm origin

A total of 26 distinct embryo:endosperm balances were identified in the seeds (Fig. 8a, b), representing a substantial increase compared to the six and seven balances reported from hand-pollinated crosses of octoploid and heptaploid mothers, respectively (Da Luz-Graña et al., 2025). Similar results have been reported in other apomictic species complexes or studies comparing multiple species within a genus (Krahulcová *et al*., 2004; Talent and Dickinson, 2007; Šarhanová *et al*., 2012; Dobeš *et al*., 2013b; Lepší *et al*., 2019). Several of the observed combinations suggest that deviations from the 2:1 maternal: paternal genome contribution to the endosperm do not result in deleterious effects (as reviewed by Vinkenoog *et al*. (2003)). This pattern is also observed in other polyploids such as *Rubus* (Šarhanová *et al*., 2012), *Hypericum* (Qu *et al*., 2010), *Potentilla* (Dobeš *et al*., 2013b) and *Crataegus* (Talent and Dickinson, 2007). In these cases, it has been postulated that the mechanisms associated with imprinting, which typically regulate endosperm development, may be either less critical or operate differently (Vinkenoog *et al*., 2003). This is advantageous in pseudogamous apomictic plants, where pollination is essential for seed formation (Asker and Jerling, 1992; Koltunow, 1993).

The 5C endosperm (2:5 balance) is the most frequent in the studied populations (Fig. 8a), typical of diplosporic apomictic pseudogamous seeds where a reduced spermatid fertilizes two polar nuclei of an unreduced ES (Matzk *et al*., 2000). For 6C endosperm (2:6 balance), we propose that it likely originates from double fertilization of the central cell, rather than fertilization by an unreduced spermatid. Double fertilization has been widely documented in pseudogamous apomictic species (Talent and Dickinson, 2007; Qu *et al*., 2010; Šarhanová *et al*., 2012; Dobeš *et al*., 2013b; Lepší *et al*., 2019). In contrast, unreduced male gametes appear to be rare, as indicated by their infrequent occurrence in the remaining seeds analysed in this study. Similar interpretations were made in *Sorbus* seeds and then corroborated by FCSS of pollen (Lepší *et al*., 2019). For the 5C+6C endosperm (2:5:6 balance), we propose that a second fertilization event occurs after the divisions of the first product of fertilization have begun. The post-zygotic fusion of a second sperm cell with a central cell that has already undergone nuclear fusion and early divisions has been previously proposed for sexually derived seeds, resulting in endosperm with different ploidy levels, named mosaic endosperm (Maheshwari, 1950; Rutishauser, 1982). The term *mosaic* endosperm originates from the observation of endosperms exhibiting phenotypic differences—such as variation in pigmentation and storage content—within individual seeds. Such cases were first reported in species such as *Zea mays*, *Petunia sp.*, *Lycopersicon esculentum*, and *Acorus calamus* (Weber, 1900; Ferguson, 1927; Bhaduri, 1933; Buell, 1938, as cited in Maheshwari, 1950) and later confirmed through cytological studies in *Ranunculus auricomus* (Rutishauser and la Cour, 1956, as cited in Rutishauser, 1982). Mosaic endosperms in *Psidium cattleyanum* f. *lucidum* provide evidence of both the involvement of two spermatids in endosperm formation and the presence of more than two polar nuclei in the central cell. The latter has already been reported in diplosporic ES (Nogler, 1984; Talent and Dickinson, 2007; Souza and Speroni, 2017; Da Luz-Graña *et al*., 2025), and it was observed in the 2:3:5 balances (and other rare cases, Fig. 8a y b), where the 3C endosperm originates from the fusion of a single unreduced polar nucleus with one reduced spermatid. Furthermore, formation of an extra autonomous endosperm from the third polar nucleus (3:5:2; 4:6:2) was also observed (Figure 8b). Unexpectedly, a third polar nucleus was also observed in reduced ES (balances 1:3:5, 1:3:4, and 1:3:2; Fig. 8a). While this feature is commonly associated with diplospory, embryo sacs with supernumerary polar nuclei have also been reported in sexual species such as *P. guajava* (Narayanaswami and Roy, 1960).

Notably, several of the above mentioned traits have been seen in other apomictic species, such as the simple fusion of a spermatid with a polar nucleus in *Crataegus* and *Potentilla* (Talent and Dickinson, 2007; Dobeš *et al*., 2013b), the formation of autonomous endosperms in pseudogamous tetraploid *Rubus* (Šarhanová *et al*., 2012) and the delayed fusion of the second sperm cell to the central cell in *Crataegus* (Talent and Dickinson, 2007). However, none of these cases have reported mosaic endosperms. Only a single endosperm tissue is interpreted from the histogram peaks, while the remaining peaks are considered endoreduplication. Here, based on our knowledge of gametophyte ontogeny and the consistency of these results and previous work in the species (Da Luz-Graña *et al*., 2025), we propose that multiple endosperm peaks may reflect true mosaic endosperm formation in *Psidium cattleyanum* f. *lucidum*.

### Embryo:endosperm balances among populations

The pentaploid LT-5x population exhibited the most heterogeneous seed progeny, resulting in the formation of 18 distinct embryo:endosperm balance types (Figures 8a, b). Seed development in this population involved contributions from both reduced and unreduced ES, as well as the frequent presence of three nuclei within the central cell (Table 4). Notably, this population also showed the highest proportion of unreduced sperm cells (15%), compared to values ranging from 1% to 5% in the other populations (Table 4). The heptaploid R13-7x, octoploid BA-8x, and hexaploid LC-6x populations also produced seeds via both ES types and consistently presented three nuclei in the central cell. These populations, however, exhibited only about half the number of genomic balance combinations observed in LT-5x. In contrast, the hexaploid CM-6x population exclusively formed seeds from unreduced ES, with minimal involvement of a third nucleus in the central cell, yielding only six distinct embryo:endosperm types.

These findings suggest that variation in the types of male and female gametes involved in seed formation—whether of reduced or unreduced—, the number of nuclei present in the central cell and the ploidy levels seem to influence the reproductive strategy of each population. Previous studies examining different taxa and ploidy levels within apomictic species complexes have reported general trends linking ploidy with reproductive mode. In most cases, diploids are predominantly sexual, triploids are largely apomictic, and tetraploids tend to be mainly apomictic, although the proportion of reproductive modes varies considerably among taxa (Krahulcová *et al*., 2004; Šarhanová *et al*., 2012; Mraž and Zang, 2019). However, comprehensive analyses including higher ploidy levels remain scarce. Notable exceptions include studies on *Potentilla puberula* (Dobeš *et al*., 2013b) and *Sorbus* (Lepší *et al*., 2019) and now *Psidium cattleyanum,* which exhibits multiple ploidy levels and distinct reproductive strategies associated with each.

### Reproductive strategy and genetic diversity within populations

The tetraploid population, ICAR-4x, used as a control in the FCSS analyses, exhibited predominantly sexual reproduction (99%), corroborating earlier suggestions of sexuality in tetraploid populations of *Psidium cattleyanum* (Machado *et al*., 2022). This pattern resembles that observed in other apomictic polyploid taxa, where diploid individuals typically reproduce sexually, while polyploids predominantly exhibit apomictic reproduction (see references in the Introduction). In certain polyploid complexes where the diploid cytotype is absent or unknown, as in *P. cattleyanum,* tetraploids have been postulated as functional diploids (e.g., *Potentilleae puberula*; Dobeš *et al*., 2013b, and *Hieracium*; Mráz and Zdvořák, 2019), and this may also apply to *P. cattleyanum* as well, as proposed by Machado *et al*. (2022).

Despite apomixis being the predominant reproductive strategy in most studied populations, the observed genetic diversity in Uruguayan populations is relatively high. Similar observations were made in *Miconia albicans*, an obligate apomictic species, where the genetic diversity rates were evaluated with ISSRs (Dias *et al*., 2018). In this species, pollen is sterile, but diplospory is of the taraxacum type (Nogler, 1984), and thus, diversity arises from recombination in restitutive meiosis during megaspore mother cell formation (Dias *et al*., 2018). In *Psidium cattleyanum*, the formation of both female and male gametes, which can be reduced, unreduced, and aneuploid, also contributes to genetic variability, through meiotic recombination and even chromosome transfer via cytomixis, as observed during microsporogenesis in this species (Souza-Pérez *et al*., 2021). Additionally, in facultative apomictic species, occasional sexual events can act as a source of genetic variation among clones (Hörandl, 2024).

The pentaploid population (LT-5x) exhibited an intermediate frequency of sexual reproduction relative to the other populations. Its values did not differ significantly from those of the hexaploid populations, which are predominantly apomictic, nor from the octoploid populations, which showed higher frequencies of sexual reproduction. Reports from other taxa show variable reproductive patterns in pentaploids: in *Potentilla puberula*, pentaploids are obligate apomicts (Dobeš *et al*., 2013b), whereas in *Sorbus*, approximately half of the pentaploids reproduce via obligate apomixis and the other half reproduce sexually (Lepší *et al*., 2019). In terms of genetic diversity, LT-5x ranked among the three most diverse populations, along with LC-6x and BA-8x (Table 2). Odd ploidy cytotypes can play an important role as mediators of gene flow (Kolář *et al*., 2017). As mentioned earlier, this population with an odd ploidy level produced the most heterogeneous seed progeny regarding ploidy levels (Figs 3 and 7) and exhibited the greatest number of distinct embryo:endosperm balance seed types (Figs 8a, b).

The two hexaploid populations (CM-6x and LC-6x) exhibited the lowest frequency of sexual reproduction, with values around 7%. In comparison, hexaploid accessions in other taxa have shown slightly higher rates of sexual reproduction, such as 16% in *Potentilla puberula* (Dobeš *et al*., 2013b) and 9.1% in *Pilosella rubra* (Doležal *et al*., 2021). Marked differences were also observed between CM-6x and LC-6x in terms of genetic diversity. LC-6x displayed the highest diversity indices among all populations, characterized by the presence of multiple embryo:endosperm balance types and a substantial proportion of reduced embryo sacs contributing to sexually derived embryos. In contrast, CM-6x showed intermediate diversity values, fewer seed types, and 100% unreduced ES.

The heptaploid and octoploid populations exhibited the highest percentages of sexuality, with R13-7x showing 33% and BA-8x showing 22%. Comparable reports for these ploidy levels include studies on cultivated octoploid *P. c.* f*. lucidum* and heptaploid P*. c.* f*. cattleyanum*, where the frequency of sexual reproduction in hand-pollinations was reported as 23% in the former and 3% in the latter (Da Luz-Graña et al., 2025). Regarding the genetic diversity rates, the populations display contrasting data. While the octoploid population (BA-8x) shows high genetic diversity, the heptaploid population (R13-7x) exhibits the lowest values among all populations. The reproductive strategy characteristics between these populations are similar, and surprisingly, R13-7x exhibits the most heterogeneous ploidy level progeny of both.

The differences in genetic diversity among populations do not directly correspond to the observed differences in sexual reproduction frequencies. It is important to consider that current patterns of genetic diversity reflect the cumulative outcome of multiple reproductive events over successive generations, whereas the reported sexual reproduction frequencies are based on a single reproductive cycle. Moreover, genetic diversity in apomictic populations is influenced by origin, demographic history, and gene flow (Kolář *et al*., 2017; Baduel *et al*., 2018; Meirmans *et al*., 2018). The reproductive strategy may fluctuate with the inclusion of data from additional years. For instance, in perennial species, the proportion of asexual versus sexual seed formation has been shown to vary across years and generations, as documented in the alpine species *Ranunculus kuepferi* (Klatt *et al*., 2018).

Remarkably, molecular variance analysis (AMOVA) and DAPC revealed strong genetic structure in the Uruguayan populations of *Psidium cattleyanum*, indicating low gene flow among them. Machado *et al*. (2022) suggested that the genetic structure of Brazilian populations is strongly influenced by geographic distance and the presence of cytotypes. Our data, however, reveal that the genetic structure is strongly influenced by the mode of reproduction (ϕCT = 0.58; p < 0.001), with minimal influence attributed to cytotypes (ϕCT = –0.23; p = 0.995), indicating that incorporating knowledge of reproductive modes could improve the understanding of the species. The prevalence of apomixis as a reproductive mode facilitates the accumulation of genetic differences among populations. Once established, these populations appear to remain reproductively isolated, with little to no gene flow occurring among them. Therefore, in populations of *P. c.* f. *lucidum*, apomixis plays a role in preserving the maternal genotype (Cruz *et al*., 1998), whereas sexual reproduction, together with previously described reproductive events, gradually introduces genetic variation. This dynamic interaction shapes the genetic structure of each population according to its predominant reproductive strategy.

### Mother and seedling ploidy levels

The predominance of a single ploidy level per population of adult plants aligns with observations reported for many *Psidium cattleyanum* populations studied in Brazil (Machado *et al*., 2021, 2022) and with population studies of other apomictic species, where geographic separation between different ploidy levels is typically observed (e.g. *Crataegus suksdorfii,* Lo *et al*., 2009; *Ranunculus auricomus,* Karbstein *et al*., 2021*, Ranunculus kuepferi,* Ladinig *et al*., 2024). Although *P. cattleyanum* is capable of vegetative propagation via root suckering, which could contribute to explaining the ploidy homogeneity (Global Database of Invasive Species, 2025; Machado *et al*., 2021), our findings suggest that this homogeneity is primarily driven by the predominance of clonal reproduction at both the embryo and seedling levels. Nevertheless, all populations produced progeny with novel ploidy levels, a trend consistent with cytotype analyses of progenies from Brazilian wild populations (Machado and Forni-Martins, 2022). In our study, ploidy diversity was greater among embryos than among germinated seedlings (see Fig. 3) and considerably lower in adult plants. This reduction reflects strong selection pressures during germination and early developmental stages, a pattern also observed in other apomictic species such as *Hieracium rubrum* (Krahulec *et al*., 2006) and *Pilosella* (Krahulcová *et al*., 2014). One proposed mechanism underlying this reduction involves delayed expression of the “endosperm balance number” block effect (deviations from the 2 maternal:1 paternal genomic ratio), which can inhibit germination (Talent and Dickinson, 2007). In this case, however, the effect appears to act only in a subset of the seeds formed. Additionally, Machado *et al*. (2022) reported that populations of *P. cattleyanum* with higher ploidy levels tend to be restricted to extreme environmental conditions. We propose that the absence of novel cytotypes among adult plants may result from a combination of intrinsic developmental constraints, low frequency of viable variants, and unfavorable environmental selection pressures.

### Facultative apomixis and polyploidy in Psidium cattleyanum

*Psidium cattleyanum* provides evidence supporting the widely recognized relationship between polyploidy and apomictic reproductive modes in angiosperms (Asker and Jerling, 1992; Hörandl *et al*., 2024; León-Martínez & Vielle-Calzada, 2019; Mráz and Zdvorak, 2019). Across the five studied populations, various reproductive processes, such as the formation of unreduced gametes (particularly ES, typically associated with diplospory), parthenogenesis, and sexuality (often linked to meiosis), occurred in varying proportions. While the number of progenies with ploidy levels differing from that of the maternal plant is relatively low, these cases, according to the literature, reveal an overlap between distinct reproductive pathways (Koltunow and Grossniklaus, 2003), potentially serving as a source of new polyploid lineages. Several studies have shown that genes controlling diplospory and parthenogenesis are regulated independently in angiosperms (Cornaro *et al*., 2023). In apomictic plants, the sexual pathway often remains functional, and apomixis results from heterochronic gene expression that redirects cell fate during reproductive development (Cornaro *et al*., 2023).

The broad progeny variation observed in *Psidium cattleyanum* f. *lucidum* is likely a consequence of the independent regulation of apomeiosis and parthenogenesis, alongside the activation of sexuality (Nogler, 1984; Koltunow and Grossniklaus, 2003; Cornaro *et al*., 2022). Populations with higher sexual reproduction rates (R13-7x and BA-8x) also showed increased frequencies of BIII hybrids (3:5 and 4:6 genome ratios). In these ES, diplospory is initiated, but fertilization precedes or replaces parthenogenesis. On the contrary, some populations produce haploid embryos via parthenogenesis from meiotically reduced egg cells. These events represent potential pathways contributing to polyploid formation in *P. c.* f. *lucidum*.

## Conclusion

This study highlights the complexity of seed formation pathways underlying the reproductive mode of a polyploid apomictic species with multiple ploidy levels. We found that the predominant reproductive mode in natural populations of *Psidium cattleyanum* f. *lucidum* is facultative pseudogamous diplosporous apomixis. Populations with different ploidy levels exhibited varying frequencies of sexual reproduction. Moreover, we observed inter-populational differences in the frequencies of gamete types (reduced vs unreduced) produced and their combinations during seed formation, with odd-ploidy populations showing the greatest heterogeneity. Populations are highly genetically structured, and our results indicate that this pattern is primarily explained by their reproductive mode. Despite this, genetic diversity remains high, and the combination of reproductive strategy and isolation suggests that each population represents a well-preserved germplasm source. These findings are encouraging for breeding programs, as natural populations constitute valuable reservoirs of genetic material for both species- and genus-level improvement.

## Acknowledgments

M.S-P. acknowledges the Postgraduate Program at the Facultad de Agronomía, Universidad de la República, where she is pursuing her PhD studies. Further expresses her gratitude to M. Bonifacino and Cristina Trujillo (UdelaR) for their collaboration on the field trip and data collection, and Yolanda Verdún (IHSM) for her invaluable help on SSR amplifications.

## Author contributions

MSP, IH, and GS conceptualized the study. MSP and GS conducted the field data collection. MSP carried out the FCSS and SSR amplifications, while MSP and MV performed the FCSS analysis. RM and MSP conducted SSR and genetic diversity analyses. A.B. performed the statistical analysis. The manuscript was written by MSP, RM, and GS, with all authors contributing to the critical review of the results and the final version of the manuscript.

