## supplementary information for "Apomixis frequency in polyploid wild populations of *Psidium cattleyanum* f. *lucidum* (Myrteae, Myrtaceae)"

### Material and methods supplementary information
**SSR tested**

**Table 1S: SSR primers used for the detection of polymorphisms in the mother plants of Psidium cattleyanum f. lucidum. Amplification quality √ amplified, little polymorphism and peak sharpness, √√ amplified and show polymorphism, √√ amplified and show several polymorphisms and peak sharpness**

| SSR name | Repeated motif | sequence | Sequence 5´-3´ | Amplification quality | TA°C | |
|---|---|---|---|---|---|---|
| mPgCIR05 | | F | GCCTTTGAACCACATC | √ | 54 | Risterucci et al., 2005; Guavamap, 2008 |
| | | R | TCAATACGAGAGGCAATA | | | Risterucci et al., 2005; Guavamap, 2008 |
| mPgCIR09 | (GA)19 | F | GCGTGTCGTATTGTTTC | √ | 55 | Risterucci et al., 2005; Guavamap, 2008 |
| | | R | ATTTTCTTCTGCCTTGTC | | | Risterucci et al., 2005; Guavamap, 2008 |
| mPgCIR10 | (CT)12 | F | GTTGGCTCTTATTTTGGT | √ | 55 | Risterucci et al., 2005; Guavamap, 2008 |
| | | R | GCCCCATATCTAGGAAG | | | Risterucci et al., 2005; Guavamap, 2008 |
| mPgCIR11 | $(CT)_{17}$ | F | TGAAAGACAACAAACGAG | √√√ | 55 | Risterucci et al., 2005; Guavamap, 2008 |

| | | | | | | |
|---|---|---|---|---|---|---|
| | | R | TTACACCCACCTAAATAAGA | | | Risterucci et al., 2005; Guavamap, 2008 |
| mPgCIR13 | (AC)12(AT)4G(GA)2 | F | CCTTTTTCCCGACCATTACA | √√ | 55 | Risterucci et al., 2005; Guavamap, 2008 |
| | | R | TCGCACTGAGATTTTGTGCT | | | Risterucci et al., 2005; Guavamap, 2008 |
| mPgCIR16 | (TC)25 | F | AATACCAGCAACACCAA | √√ | 55 | Risterucci et al., 2005; Guavamap, 2008 |
| | | R | CATCCGTCTCTAAACCTC | | | Risterucci et al., 2005; Guavamap, 2008 |
| mPgCIR21 | (AG)15GG(AG)7 | F | TGCCCTTCTAAGTATAACAG | √√ | 55 | Risterucci et al., 2005; Guavamap, 2008 |
| | | R | AGCTACAAACCTTCCTAAA | | | Risterucci et al., 2005; Guavamap, 2008 |
| mPgCIR23 | (TA)4(GT)7 | F | GTCTATACCTAATGCTCTGG | √ | 55 | Risterucci et al., 2005; Guavamap, 2008 |
| | | R | CCCAGGAAAATCTATCAC | | | Risterucci et al., 2005; Guavamap, 2008 |
| mPgCIR26 | (GT)2(GA)17 | F | CTACCAAGGAGATAGCAAG | √√ | 55 | Risterucci et al., 2005; Guavamap, 2008 |

| | | | | | | |
|---|---|---|---|---|---|---|
| | | R | GAAATGGAGACTTTGGAG | | | Risterucci et al., 2005; Guavamap, 2008 |
| mPgCIR44 | | F | TTCCAGGTCTATTGGATGTC | √ | | Risterucci et al., 2005; Guavamap, 2008 |
| | | R | GGGGACACAAAACTTCATTA | | | Risterucci et al., 2005; Guavamap, 2008 |
| mPgCIR46 | (GA)36 | F | ATAGAACGCCATGTTACCAA | √√ | 56 | Risterucci et al., 2005; Guavamap, 2008 |
| | | R | CAGGCTTATCTGTTACACCA | | | Risterucci et al., 2005; Guavamap, 2008 |
| mPgCIR94 | (GA)18/(GT)6 | F | CAACCTTCCCGTGATTATT | √√ | 50 | Risterucci et al., 2005; Guavamap, 2008 |
| | | R | CTAGCTTCTTCAGTGGGAAC | | | Risterucci et al., 2005; Guavamap, 2008 |
| mPgCIR180 | | F | CATGGATTCAACTCTTGTCG | √ | | Risterucci et al., 2005; Guavamap, 2008 |
| | | R | CTACATTGGAAGCAGAATGG | | | Risterucci et al., 2005; Guavamap, 2008 |
| mPgCIR192 | (GA)23 | F | ACGCTAACTATCGAAATGCT | √ | 50 | Risterucci et al., 2005; Guavamap, 2008 |

| | | | | | | |
|---|---|---|---|---|---|---|
| | | R | ACTACGCACTTGATGGAGAT | | | Risterucci et al., 2005; Guavamap, 2008 |
| mPgCIR179 | (GA)16 | F | GGGTCTCGACTAAAGAAGGA | √√√ | 54 | Risterucci et al., 2005; Guavamap, 2008 |
| | | R | CCTCCATTTGCATCAACTTT | | | Risterucci et al., 2005; Guavamap, 2008 |
| mPgCIR220 | (GT)8/(GA)20 | F | AGAGCAGTGGTTGCTATTTT | √√√ | 55 | Risterucci et al., 2005; Guavamap, 2008 |
| | | R | CCCATCTCTTACTTTTCTTGTG | | | Risterucci et al., 2005; Guavamap, 2008 |
| Pca-UNICAMP01 | (CT)14(CT)8 | F | *GACTTGACAAGGGCAAAGTC | √√ | 55 | (Machado et al., 2021) |
| | | R | TAAAGGTGCATTTGTCTGCG | | | (Machado et al., 2021) |
| Pca-UNICAMP02 | (TG)9 | F | *AAGTTGGCAGGTCTAGTTCC | √ | 60 | (Machado et al., 2021) |
| | | R | TCAAGCTAGGTATGCTTCCC | | | (Machado et al., 2021) |
| Pca-UNICAMP04 | (CT)22 | F | *CCTTTTACACATTAGCTCTCTC | √√√ | 55 | (Machado et al., 2021) |
| | | R | GACCTGGGGTGTCATAACAA | | | (Machado et al., 2021) |
| Pca-UNICAMP05 | (CT)16(CA)10 | F | *CAAAGTAGGTATGCTGCGTG | √√ | 63 | (Machado et al., 2021) |
| | | R | GCAAGTTAAACCGATCTGCA | | | (Machado et al., 2021) |
| Pca-UNICAMP06 | (CT)16(CT)8 | F | *GACTTGACAAGGGCAAAGTC | √√ | 60 | (Machado et al., 2021) |
| | | R | CTGCGTGTGCTAGACCTTAA | | | (Machado et al., 2021) |

| | | | | | | |
|---|---|---|---|---|---|---|
| Pca-UNICAMP07 | (GT)8(CT)20 | F | *ACTAATGACGGTCCTTGAGAC | √ | 51 | (Machado et al., 2021) |
| | | R | TTGTTGAGACTGCATGCATG | | | (Machado et al., 2021) |
| Pca-UNICAMP08 | (AG)31(CT)26(TC)16(TG)9 | F | *GCACGTGCAAGAAAGAGAG | √√ | 60 | (Machado et al., 2021) |
| | | R | GTTCACACAGCACGCTAATT | | | (Machado et al., 2021) |
| Pca-UNICAMP09 | (CT)15(AG)23 | F | *CATGAAAAATGAGTAGGCTCTC | √ | 60 | (Machado et al., 2021) |
| | | R | CTCAGCTGGTTGTGCATAAC | | | (Machado et al., 2021) |
| Pca-UNICAMP10 | (TC)15(AC)11(CT)21 | F | *ACAACCCTTCTTTGCCCTAA | √√√ | 60 | (Machado et al., 2021) |
| | | R | ACAGATGTCATCAGAAGACACT | | | (Machado et al., 2021) |
| Pca-UNICAMP11 | (CT)11(TTC)8 | F | *CGTTATCTCCTTCCTCCGAG | √√√ | 60 | (Machado et al., 2021) |
| | | R | ATCGCCGATCAACTTCGAG | | | (Machado et al., 2021) |

**Results supplementary information**
**Statistical análisis**

**Table 2S:** Tukey test for comparison of means of apomixis within ploidy levels. Intervals are back-transformed from the logit scale. Confidence level used: 0.95, significance level used: alpha = 0.05

| Ploidy level | prob | SE | Asymp.LCL | asymp.UCL | group |
|---|---|---|---|---|---|
| 7x | 0.676 | 0.0401 | 0.593 | 0.750 | A |
| 8x | 0.791 | 0.0329 | 0.719 | 0.848 | AB |
| 5x | 0.851 | 0.0287 | 0.785 | 0.899 | B |
| 6x | 0.939 | 0.0138 | 0.906 | 0.961 | C |

**Tables 3S.**
Table 3.1S. Contingency tables with populations as rows and the frequency of each category as columns. Ploidy (2C embryos, 3C and 4Cembryos).

| Mother ploidy | 2C | 3C | 4C | Total |
|---|---|---|---|---|
| 5x | 5 | 12 | 6 | 23 |
| 6x | 5 | 12 | 1 | 18 |
| 7x | 6 | 37 | 1 | 44 |
| 8x | 5 | 27 | 0 | 32 |
| Total | 21 | 88 | 8 | 117 |

Table 3.2S. Global test of independence performed using Pearson's chi-square test and the multivariate $G^2$ (MV-G2) test.

| Statistics | Value | fd | p |
|---|---|---|---|
| Pearson Chi Square | 20.36 | 6 | 0.0024 |
| Chi Square MV-G2 | 17.84 | 6 | 0.0066 |

Tables 3.3S and 3.5S Contingency tables for pairwise comparison. Only significant differences are shown. 3.4S and 3.6S Global test of independence performed using Pearson's chi-square test and the multivariate $G^2$ (MV-G2) test. Ploidy (2C embryos, 3C and 4Cembryos)

Table 3.3S

| Mother Ploidy | 2C | 3C | 4C | Total |
|---|---|---|---|---|
| 5x | 5 | 12 | 6 | 23 |
| 7x | 6 | 37 | 1 | 44 |
| Total | 11 | 49 | 7 | 67 |

Table 3.4S

| Statistics | Value | fd | p |
| --- | --- | --- | --- |
| Pearson Chi Square | 10.91 | 2 | 0.0043 |
| Chi Square MV-G2 | 10.73 | 2 | 0.0047 |

Table 3.5S

| Mother Ploidy | 2C | 3C | 4C | Total |
| --- | --- | --- | --- | --- |
| 5x | 5 | 12 | 6 | 23 |
| 8x | 5 | 27 | 0 | 32 |
| Total | 10 | 39 | 6 | 55 |

Table 3.6S

| Statistics | Value | fd | p |
| --- | --- | --- | --- |
| Pearson Chi Square | 10.58 | 2 | 0.0050 |
| Chi Square MV-G2 | 12.76 | 2 | 0.0017 |

**Results supplementary information**

**Molecular analysis of SSR**

Table 4S- Description of the SSR markers for Uruguayan populations of Psidium cattleyanum: NM number of bands, PIC polymorphism content, DP discriminatory power, and exclusive bands for each population

| Locus' name | NB | PIC | DP | Exclusive bands | | | | |
|---|---|---|---|---|---|---|---|---|
| | | | | LT-5x | BA-8x | CM-6x | R13-7x | LC-6x |
| Pca-UNICAMP04 | 11 | 0,85 | 0,84 | 0 | 0 | 1 | 3 | 2 |
| Pca-UNICAMP10 | 29 | 0,90 | 0,85 | 3 | 1 | 1 | 1 | 12 |
| Pca-UNICAMP11 | 16 | 0,88 | 0,84 | 1 | 3 | 0 | 3 | 1 |
| mPgCIR179 | 15 | 0,86 | 0,86 | 0 | 0 | 4 | 0 | 2 |
| mPgCIR220 | 18 | 0,86 | 0,87 | 4 | 2 | 3 | 0 | 1 |
| mPgCIR11 | 13 | 0,81 | 0,85 | 0 | 0 | 0 | 0 | 8 |
| Global | | | | 8 | 6 | 9 | 7 | 26 |

Figure 1S. Psidium cattleyanum f. lucidum clusters onto a heatmap based on Bruvo's distance.

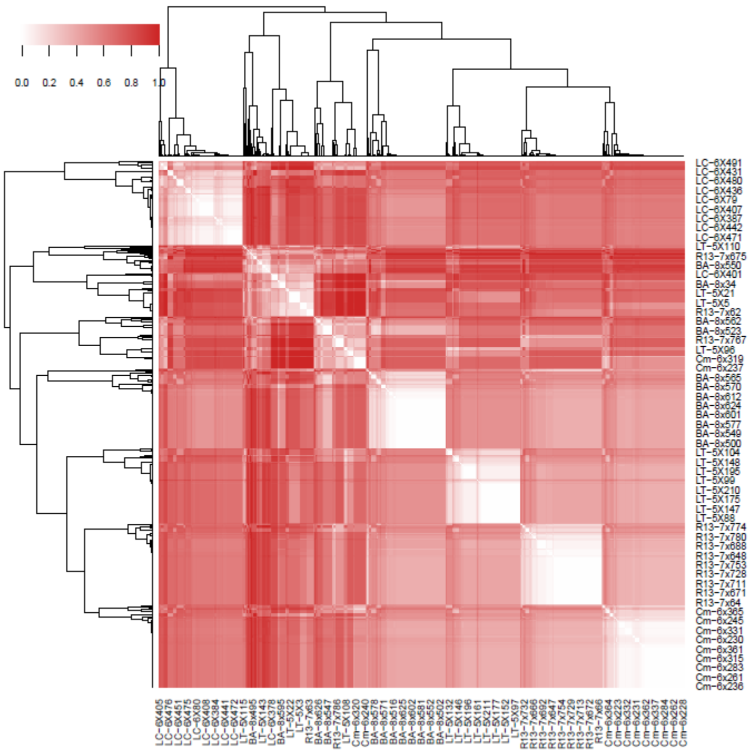
